# An Experimental Platform for Stochastic Analyses of Single Serotonergic Fibers in the Mouse Brain

**DOI:** 10.1101/2023.06.16.545362

**Authors:** Kasie C. Mays, Justin H. Haiman, Skirmantas Janušonis

**Affiliations:** Department of Psychological and Brain Sciences University of California, Santa Barbara, CA 93106-9660, USA

**Keywords:** 5-hydroxytryptamine (5-HT), serotonin, axon, varicosities, Brainbow, tortuosity, stochastic, random walk

## Abstract

The self-organization of the serotonergic matrix, a massive axon meshwork in all vertebrate brains, is driven by the structural and dynamical properties of its constitutive elements. Each of these elements, a single serotonergic axon (fiber), has a unique trajectory and can be supported by a soma that executes one of many available transcriptional programs. It necessitates the development of specialized methods for single-fiber analyses, both at the experimental and theoretical levels. We developed an integrated system that facilitates experimental isolation of single serotonergic fibers in brain tissue, including regions with high fiber densities, and demonstrated the potential of their quantitative analyses based on stochastic modeling. Single fibers were visualized using two transgenic mouse models, one of which is the first implementation of the Brainbow toolbox in this system. The trajectories of serotonergic fibers were automatically traced in the three spatial dimensions with a novel algorithm, and their properties were captured with a single parameter associated with the directional von Mises-Fisher probability distribution. The system represents an end-to-end workflow that can be imported into various studies, including those investigating serotonergic dysfunction in brain disorders. It also supports new research directions inspired by single-fiber analyses in the serotonergic matrix, including supercomputing simulations and modeling in physics.

## 1. INTRODUCTION

The structural and dynamical properties of single serotonergic fibers determine their population-scale behavior and ultimately lead to the emergence of the serotonergic matrix in all vertebrate brains. This matrix, composed of a very large number of meandering fibers and their branches, is invariably present in mammals (Wilson and Molliver, 1991; Awasthi et al., 2021), birds (Sako et al., 1986; Challet et al., 1996), reptiles (Bennis et al., 1990; Challet et al., 1991), amphibians (Ueda et al., 1984; Bhat and Ganesh, 2023), bony fish (Lillesaar, 2011; López and González, 2014), and cartilaginous fish (Stuesse et al., 1995; Carrera et al., 2008). The fundamental role of the serotonergic matrix remains enigmatic; in adulthood, it may support neuroplasticity (Lesch and Waider, 2012; Teissier et al., 2017; Huang et al., 2021; Daws et al., 2022; Morgan et al., 2023), with potential implications for artificial neural networks (Lee et al., 2022).

The serotonergic fibers are long axons of neurons that express the tryptophan hydroxylase 2 gene (*Trp2*) and can synthesize the neurotransmitter serotonin (5-hydroxytryptamine, 5-HT) from the amino acid tryptophan. In mammals, their somata are restricted to the raphe nuclei of the brainstem. The rostral raphe nuclei (the dorsal raphe, the median raphe, and the caudal linear nucleus) project anteriorly to supply fibers to virtually all midbrain and forebrain brain regions (Jacobs and Azmitia, 1992; Hornung, 2003).

The deeper structure of this system, as revealed by recent studies, may require novel conceptual frameworks. Serotonin is not the only neuroactive compound that can be produced from tryptophan (*e.g.*, it can also be metabolized into kynurenic acid, quinolinic acid, or *N*,*N’*-dimethyltryptamine (DMT)) (Dean et al., 2019; Yabut et al., 2019; Paul et al., 2022; Vargas et al., 2023), many serotonergic neurons also release glutamate and other neurotransmitters (and therefore can be considered “other-ergic” neurons that co-release serotonin) (Okaty et al., 2019), and the population of the serotonergic neurons is now known to consist of transcriptionally diverse, nested cell clusters (Okaty et al., 2019; Ren et al., 2019; Okaty et al., 2020), some with distinct neuroanatomical profiles (Ren et al., 2018; Okaty et al., 2020).

The transcriptomics studies have highlighted the importance of the understanding of serotonergic neurons at the single-cell level (Okaty et al., 2020). Individual neurons can choose among several “adjacent” transcriptional networks, depending on their developmental path and activation state (Okaty et al., 2019), and are also affected by stochastic noise (Elowitz et al., 2002; Raj and van Oudenaarden, 2008). These networks, as detected in actual neurons, are likely to represent dynamical attractors and are therefore computationally constrained (*i.e.*, not every set of gene activity is equally stable).

Further, individual serotonergic fibers acquire unique identities because their trajectories are strongly stochastic (Janušonis and Detering, 2019). It does not mean that any trajectory is possible in practice. Individual trajectories may represent “sample paths” (realizations) of rigorously-definable spatial stochastic processes, which again implies computational constrains. We have demonstrated the potential of this approach in recent studies in which serotonergic axons were modeled as paths of reflected fractional Brownian motion (FBM), a stochastic continuous-time process (Janušonis et al., 2020; Janušonis et al., 2023). In these supercomputing simulations, performed in geometric shapes based on the mouse brain, the resultant fiber densities approximated the actual serotonergic fiber densities, with no other biological information. These models require improvements to reflect neural tissue heterogeneities (Wang et al., 2023), but they can eventually bridge the stochasticity on the microscopic scale (the uniqueness of fiber trajectories in each brain) and the determinism on the macroscopic scale (the predictability of regional fiber densities (Awasthi et al., 2021)).

The serotonergic neurons are among the first neurons to acquire a mature phenotype in the developing brain (Lidov and Molliver, 1982; Hendricks et al., 1999; Hawthorne et al., 2010). Somewhat paradoxically, they also appear to remain highly dynamic in the adult brain. In particular, serotonergic fibers can robustly regenerate after a traumatic injury (Jin et al., 2016; Kajstura et al., 2018; Cooke et al., 2022), with new paths (Jin et al., 2016). This potential and the challenge of maintaining the physical continuity of extremely long, unfasciculated fibers suggests that a healthy adult brain may always contain degenerating and regenerating serotonergic fibers. This in turn implies that the fiber trajectories of given serotonergic neurons may change during an individual’s life span. Because of technical challenges, this prediction currently cannot be directly verified in experimental studies. However, it is known that the maintenance of the serotonergic fiber matrix depends on the *adult* expression of *Lmx1b* and *Pet1*, two genes essential for the embryonic development of serotonergic neurons (Kitt et al., 2022), and that the stochastic serotonergic trajectories acquire interesting computational properties if they do not stay constant (Lee et al., 2022).

The large-scale properties of the serotonergic matrix can be theoretically predicted from the properties of single serotonergic fibers – but not the other way around. Specifically, the same patterns of regional fiber densities can be produced by different models, and major alterations in fiber densities, such as in autism (Azmitia et al., 2011) or epilepsy (Maia et al., 2019), does not imply specific changes in the properties of single fibers. Importantly, these properties include not only trajectories, but the pattern and probability distribution of branching events, the spatial and temporal features of varicosities (fiber “swellings”), and other features. The branching of serotonergic fibers remains poorly understood because their paths often cross at distances that fall below the limit of optical resolution, making the definitive identification of branching points difficult even in three-dimensional, high-resolution images (Janušonis et al., 2019). Branching serotonergic fibers are more easily accessible in primary brainstem cultures (Hingorani et al., 2022), but the artificial culture environments limit the applicability of these observations in natural neural tissue. The field also lacks a fundamental understanding of the serotonergic varicosities. Their distribution may reflect the developmental age of the fiber (Maddaloni et al., 2017), but the same serotonergic fiber may also contain varicosities that differ in size, shape, and spacing – both in brain tissue (Baizer, 2001; Gagnon and Parent, 2014; Maddaloni et al., 2017) and *in vitro* (Hingorani et al., 2022). This variability does not necessarily imply different varicosity types and may instead reflect a dynamic process frozen at different stages along the fiber trajectory (in fixed preparations). Serotonergic varicosities are likely to be fluid structures, as suggested by experimental studies of mice and *Drosophila* with elevated serotonin levels (Daubert et al., 2010), a rat model of epilepsy (Maia et al., 2019), and analyses of varicosities in other neurotransmitter systems. Specifically, the appearance of varicosities may reflect the current state of the fiber segment, the effects of its local microenvironment, and purely stochastic fluctuations (Hellwig et al., 1994; Hatada et al., 1999; Shepherd et al., 2002; Gu, 2021; Ma et al., 2022). Notably, the dynamics of varicosities on the same fiber are likely to be correlated, necessitating unambiguous discrimination among individual fibers that often cross and intertwine in the same spatial location. An added complexity is that different serotonergic fibers in the same location may execute different transcriptional programs. Therefore, disregarding the identity of fibers is equivalent to a gross distortion of the covariance structure among these various elements, potentially leading to incorrect statistical calculations and conclusions.

At present, reliable identification and analyses of extended segments of single serotonergic fibers present serious challenges, especially in brain tissue. In particular, it requires knowledge about the continuity of a given fiber. Only a handful recent studies have focused on descriptions of single serotonergic fibers at this level of precision (Gagnon and Parent, 2014; Maddaloni et al., 2017; Janušonis et al., 2019; Hingorani et al., 2022). Furthermore, specialized quantitative methods for their analyses are virtually absent. For example, the tortuosity index, often used to describe meandering fibers (Jin et al., 2016; Pratelli et al., 2017), is a relatively unstable metric in that it depends on the measured fiber length and imaging dimensionality, and also can vary drastically in different realizations of the same spatial stochastic process.

This study aimed to (*i*) develop flexible and accessible experimental mouse models to isolate single serotonergic fibers in brain tissue, independently of their serotonin content or the expression of the serotonin transporter (SERT), (*ii*) build an algorithm for high-precision tracing of single fibers in noisy three-dimensional images, and (*iii*) develop a statistical model to achieve a succinct description of fiber trajectories that can vary strongly in their individual appearance. The three parts constitute a workflow that can be imported into various experimental setups.

## 2. MATERIALS AND METHODS

### 2.1. Transgenic Mice

Three transgenic mouse lines were used in the study: *Tph2*-*iCreER* (Tg(Tph2-icre/ERT2)6Gloss/J; RRID: IMSR_JAX:016584), *ROSA*^mT/mG^ (B6.129(Cg)-Gt(ROSA)26Sor^tm4(ACTB-tdTomato,-EGFP)Luo^/J; RRID: IMSR_JAX:007676), and *Brainbow 3.2* (line 7) (Tg(Thy1-Brainbow3.2)7Jrs/J; RRID: IMSR_JAX:021227). The *Tph2*-*iCreER* mice express a tamoxifen-inducible Cre recombinase under the control of the promoter of the tryptophan hydroxylase 2 gene (*Tph2*) that encodes a key enzyme in the serotonin-synthesis path. The *ROSA*^mT/mG^ (Cre reporter) mice ubiquitously express a cell membrane-targeting tdTomato, which after Cre-recombination is replaced with a cell membrane-targeting enhanced green fluorescent protein (EGFP) (Muzumdar et al., 2007). The *Brainbow* mice contain the *Brainbow 3.2* construct under neuron-specific regulatory elements from the *Thy1* gene (Cai et al., 2013). This construct includes sequences for three cell membrane-targeting (farnesylated) fluorescent proteins (mOrange2, EGFP, and mKate2) that after Cre-recombination are expressed in random intensity combinations in individual cells. The construct also includes a fourth sequence (in the stop cassette) for a mutated (non-fluorescent and nucleus-targeting), *Phialidium*-derived yellow fluorescent protein (PhiYFP(Y65A)) that can be used to assess the transcriptional activity of the locus in specific brain regions, before recombination (Cai et al., 2013).

The breeders were purchased from The Jackson Laboratory (JAX), and the colonies were kept in a vivarium on a 12:12 light-dark cycle with free access to food and water. The *ROSA*^mT/mG^ mice were kept and bred in the homozygous state, as suggested by the supplier. Mouse litters, including the offspring of crosses, were PCR-genotyped by using DNA from toe biopsies collected at postnatal day 7. Genomic DNA was isolated with the QIAamp Fast DNA Tissue Kit (Qiagen #51404), and the transgene sequences were amplified with an Eppendorf Mastercycler pro S using iTaq DNA polymerase (Bio-Rad #1708870), a dNTP mix (Bio-Rad #1708874), and primers with sequences provided by JAX (using the cycling protocol suggested by the supplier of the polymerase). The amplicons were analyzed with a 2200 TapeStation (Agilent Technologies). All animal procedures have been approved by the UCSB Institutional Animal Care and Use Committee.

### 2.2. EGFP Labeling of Serotonergic Fibers

*Tph2*-*iCreER* and *ROSA*^mT/mG^ mice were crossed, and their offspring with the transgenes were allowed to reach adulthood (at least 10 weeks of age). The induction of Cre-recombination was achieved by administering tamoxifen (Millipore-Sigma #T5648) dissolved in corn oil (Millipore-Sigma #C8267). The stock concentration (20 mg/mL) was stored at −20°C, and mice were injected intraperitoneally with around 0.1 mL of the solution (at 75 mg/kg) for 5 consecutive days. Following the tamoxifen treatment, the mice were allowed to survive for one week to one month.

### 2.3. Stochastic Multicolor Labeling of Serotonergic Fibers with *Brainbow* AAVs

The *Brainbow* adeno-associated viruses (AAVs) (AAV-EF1a-BbTagBY [Addgene #45185-AAV9] and AAV-EF1a-BbChT [Addgene #45186-AAV9]) (Cai et al., 2013) were stored at −75°C. Before use, they were diluted to 1.5×10^12^ vg/mL each in a tube containing sterile, alcohol-free saline. Adult (at least 10 weeks of age) *Tph2*-*iCreER* mice were anesthetized with an intraperitoneal injection of a mixture of ketamine (100 mg/kg) and xylazine (10 mg/kg) and placed in a small-animal stereotaxic frame. Further anesthesia was maintained with inhaled isoflurane, and the animal’s core temperature was maintained at 37°C using a TCAT-2DF temperature controller (a closed loop rectal probe-heating pad system; Physitemp Instruments). The incision area (with hair removed) was given a subcutaneous injection of lidocaine and disinfected with Betadine. A rostrocaudal skin incision was made with a scalpel, and a small hole was drilled in the skull directly over the DR. The needle of a 10 µL-Hamilton microsyringe was lowered into the dorsal raphe, and 1-2 µL of the AAV mixture was slowly pressure-injected at the lambda, 3.5 mm ventral to the dura. The needle remained in the DR for 5-10 minutes and was slowly withdrawn. The skin incision was closed with sterile wound clips, and the animal was monitored until it fully recovered.

One week after the surgery, the induction of Cre-recombination was achieved as described for the EGFP labeling. Following the tamoxifen treatment, the mice were allowed to survive for 1-3 months to ensure fluorophore transport to distal axon segments.

We also attempted to label serotonergic fibers by crossing *Tph2*-*iCreER* and *Brainbow* mice. Their offspring with the transgenes were allowed to reach adulthood (at least 10 weeks of age), and the induction of Cre-recombination was performed as described for the EGFP labeling. Following the tamoxifen treatment, the mice were allowed to survive for around one month.

### 2.4. Immunohistochemistry (IHC)

Mice were euthanized with CO_2_ and their brains were dissected into 0.1 M phosphate-buffered saline (PBS, pH 7.2). They were immediately immersion-fixed in 4% paraformaldehyde overnight at 4°C, cryoprotected in 30% sucrose for two days at 4°C, and sectioned coronally at 40 µm thickness on a freezing microtome. The sections were stored in PBS and processed further with immunohistochemistry.

#### 2.4.1. Cre IHC

Sections were rinsed in PBS and treated with a heat-induced epitope retrieval (HIER) procedure (untreated sections produced no signal). The HIER was performed in a basic Tris-HCl buffer solution (10 mM, pH 9.0), with 30 second-microwave pulses at 30-50% power for 4-6 minutes (to prevent section disintegration). The sections were rinsed in PBS, blocked in 2% normal donkey serum (NDS) for 30 minutes at room temperature, and incubated on a shaker in rabbit anti-Cre IgG (1:1000; Abcam #ab216262) with 2% NDS and 0.5% Triton X-100 (TX) in PBS for 3 days at 4°C. They were rinsed in PBS three times (10 minutes each) and incubated in Cy3-conjugated donkey anti-rabbit IgG (1:200; Jackson ImmunoResearch #711-165-152) with 2% NDS and 0.3% TX for 90 minutes at room temperature. The sections were rinsed in PBS three times (10 minutes each), mounted onto gelatin/chromium-subbed glass slides, allowed to air-dry, and coverslipped with ProLong Gold Antifade Mountant with no DAPI (ThermoFisher Scientific, #P36930).

#### 2.4.2. PhiYFP(Y65A) IHC

Sections were rinsed in PBS, blocked in 2% normal donkey serum (NDS) for 30 minutes at room temperature, and incubated on a shaker in rabbit anti-PhiYFP(Y65A) IgG (1:500; kindly provided by Dr. Dawen Cai) with 2% NDS and 0.5% TX in PBS for 3 days at 4°C. They were rinsed in PBS three times (10 minutes each) and incubated in Cy3-conjugated donkey anti-rabbit IgG (1:400; Jackson ImmunoResearch #711-165-152) with 2% NDS for 90 minutes at room temperature. The sections were rinsed in PBS three times (10 minutes each), mounted onto gelatin/chromium-subbed glass slides, allowed to air-dry, and coverslipped with ProLong Gold Antifade Mountant with no DAPI.

#### 2.4.3. EGFP IHC

Sections were rinsed in PBS, blocked in 2% normal donkey serum (NDS) for 30 minutes at room temperature, and incubated on a shaker in rabbit anti-GFP IgG (1:500; Abcam #ab6556) with 2% NDS and 0.3% TX in PBS for 2-3 days at 4°C. They were rinsed in PBS three times (10 minutes each) and incubated in AlexaFluor 488-conjugated donkey anti-rabbit IgG (1:1000; ThermoFisher #A21206) with 2% NDS for 90 minutes at room temperature. The sections were rinsed in PBS three times (10 minutes each), mounted onto gelatin/chromium-subbed glass slides, allowed to air-dry, and coverslipped with ProLong Gold Antifade Mountant with DAPI (ThermoFisher Scientific, #P36931).

#### 2.4.4. Brainbow IHC

Sections were rinsed in PBS, blocked in 2% normal donkey or goat serum (NDS or NGS, matching the host of the secondary antibodies) for 30 minutes at room temperature, and incubated on a shaker in rabbit anti-GFP IgG (1:500; Abcam #ab6556), rat anti-mTFP IgG (1:500; Ximbio #155264), and guinea pig anti-TagRFP IgG (1:500; Ximbio #155267), with 2% NDS or NGS and 0.3% TX in PBS for 3 days at 4°C. They were rinsed in PBS three times (10 minutes each) and incubated in either a donkey or goat set of secondary antibodies diluted in PBS with 2% NDS or NGS, respectively, for 90 minutes at room temperature. The donkey set consisted of AlexaFluor 488-conjugated donkey anti-rabbit IgG (1:1000; ThermoFisher #A21206), AlexaFluor 594-conjugated donkey anti-rat IgG (1:500; Jackson ImmunoResearch #712-585-150), and AlexaFluor 647-conjugated donkey anti-guinea pig IgG (1:500; Jackson ImmunoResearch #706-605-148). The goat set consisted of AlexaFluor 488-conjugated goat anti-rabbit IgG (1:1000; ThermoFisher #A11034), AlexaFluor 546-conjugated goat anti-rat IgG (1:1000; ThermoFisher #A11081), and AlexaFluor 647-conjugated goat anti-guinea pig IgG (1:1000; ThermoFisher #A21450). The sections were rinsed in PBS three times (10 minutes each) and stored in PBS at 4°C or immediately mounted onto gelatin/chromium-subbed glass slides, allowed to air-dry, and coverslipped with ProLong Gold Antifade Mountant with no DAPI. Initially, the primary antibody set also included chicken anti-mCherry IgY (1:500; Ximbio #155259) and the secondary antibody set included either AlexaFluor 594-conjugated donkey anti-chicken IgY (1:500; Jackson ImmunoResearch #703-585-155) or AlexaFluor 546-conjugated goat anti-chicken IgY (1:1000; ThermoFisher #A11040). However, this fourth signal was eventually eliminated because of its relative weakness and overlap with another imaging channel. A similar reduction of fluorophore channels from four to three was used by the authors of the Brainbow system (Cai et al., 2013). The two secondary antibody sets produced comparable results.

### 2.5 Microscopy Imaging

Epifluorescence imaging was performed in one or two of three channels (Cy3, GFP, and DAPI) on a Zeiss AxioVision Z1 using a 5× objective (NA 0.13) or 10× objective (NA 0.45).

Confocal imaging was performed in two channels (AlexaFluor 488 and DAPI) or three (Brainbow) channels (AlexaFluor 488, AlexaFluor 594 or 546, and AlexaFluor 647) on a Leica SP8 resonant scanning confocal system. High-power images were obtained using a 63×oil objective (NA 1.40) with the XY-resolution of 60-70 nm/pixel and the Z-resolution of 300 nm/optical section. Typical z-stacks consisted of around 100 optical sections. The figures show maximum-intensity projections.

### 2.6. Computational Analyses

In simulations, sample unit-vectors drawn from the von Mises-Fisher (vMF) directional probability distribution were generated using the Wood algorithm (Wood, 1994; Hoff, 2009). The vMF distribution is parametrized by the concentration parameter (κ > 0), which is analogous to the inverse of the variance of the normal distribution (Janušonis and Detering, 2019). Briefly, the algorithm first generates a sample vector that has the required distribution around the fixed mean direction of ***μ*^0^** = (0, 0, 1) and then rotates the vector to any other mean direction ***μ*** = (*μ*_1_, *μ*_2_, *μ*_3_) (where *μ* is again a unit-vector). In the first step, (*i*) a 2D-sample unit-vector ***ν*** is generated from the uniform distribution on the circle, (*ii*) a sample scalar *w* is generated from (−1, 1) from the probability distribution that is proportional to *e^kw^* (*e.g.*, using rejection sampling), and (*iii*) the 3D-sample vector ***u*^0^** is then constructed as 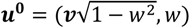. In the second step, the obtained vector is rotated to the required population mean with ***Mu*^0^**, where the orthogonal 3 × 3 matrix ***M*** can be given by

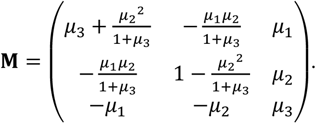

Fibers can then be simulated with repeated sampling, in which the direction of each step has the vMF distribution with a constant κ and a mean direction iteratively defined as the direction of the previous step.

Conversely, if the directions of all steps are already known (*e.g.*, from experimental data), the fiber can be iteratively rotated such that the current step direction always coincides with ***μ*^0^** = (0, 0, 1), and then the direction of the next step can be determined with respect to this standardized direction. If the original next direction is given by ***u***, the standardized next direction is given by ***M***^−1^***u***, where the elements of **M** are constructed from the original current direction ***μ***. In particular, if the next direction is the same as the current direction, one obtains ***M***^−1^***μ*** = ***μ*^0^**, as expected. After all directions have been standardized to the corresponding previous directions, this unit-vector *sample* is used to estimate κ. Several methods are available that are based on the *length* of the mean vector of the sample (*L*, where 0 ≤ *L* ≤ 1). If all steps maintain the same direction (*i.e.*, the fiber is “infinitely” rigid), *L* = 1. If, conversely, all directions are equally possible (including the unrealistic return to the previous point), *L* = 0 (in the limit). Therefore, higher *L* values should correspond to lower κ values. A precise estimate of κ for any *L* can be obtained by using a method based on a fixed-point iteration (Tanabe et al., 2007). More conveniently, a relatively accurate κ estimate can also be obtained with the formula (Tanabe et al., 2007)

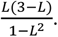

The reported κ values were calculated with this method. It should be noted that, if *L* ≥ 0.9, an even simpler formula can be used, 1/(1 − *L*) (Mardia and Jupp, 2000). It follows that this formula can be used to estimate κ values that are greater than 10 (which makes it highly applicable in studies of serotonergic fibers). Both formulas assume that *L* has been calculated in the three spatial dimensions; however, *L*-based estimators for lower and higher dimensions are available (Mardia and Jupp, 2000; Tanabe et al., 2007). The toolbox of directional statistics is rapidly expanding and includes goodness-of-fit and similarity tests, regression models, and other methods analogous to but mathematically different from those in classical statistics (Pewsey and Garcia-Portugués, 2021).

All simulation and analysis scripts were written in Wolfram Mathematica 13.

## 3. RESULTS

### 3.1. The Isolation of Single Fibers with EGFP Labeling

The induction of Cre-recombination is an inherently stochastic event at the single-cell level, but the efficiency of this process can be partially controlled with the dose and administration regimen of the inducing agent (tamoxifen) (Zhang et al., 2005; Yoshinobu et al., 2021). Once the production of the fluorophore has been initiated, it can visualize all cell processes, especially if the fluorophore targets the cell membrane (Muzumdar et al., 2007; Cai et al., 2013). These properties can support the visualization of sparse, theoretically random subsets of neurons and their axons (Economo et al., 2016), with each axon fully labeled (given a sufficient transport time).

Following the induction of the *Tph2*-dependent Cre-recombination, strong EGFP expression was observed in the rostral raphe nuclei. EGFP-positive somata were detectable in unstained sections (Fig. 1A); the signal was further verified and enhanced with GFP-immunohistochemistry (Figs. 1B, C). After a relatively short post-induction time (less than two weeks), uninterrupted single fibers could be visualized in various brain regions (Figs. 1D-K). Some of these fibers could be traced over considerable distances (Fig. 1D) and contained rich statistical information about the underlying stochastic process. These fibers were used in the random-walk modeling described further. Some fibers had growth cone-like swellings (Figs. 1E, F) and resembled developing serotonergic axons in the embryonic mouse brain (Hingorani et al., 2022). However, these structures require further analyses because their functional identity and terminal location cannot be confirmed in this preparation (*e.g.*, the fiber may continue in an adjacent section). Some fibers in several regions (*e.g.*, the inferior colliculus and the dentate gyrus) contained fibers with multiple large swellings (Figs. 1G, H). They resembled spheroids, abnormal varicosities associated with dystrophic serotonergic fibers (Daubert et al., 2010). These segments may be in the process of natural degeneration, also because their general morphology resembles that of collapsing axons disconnected from their soma (Shaw and Bray, 1977; Baas and Heidemann, 1986), but they may also represent fibers making extension or branching decisions. True branching events were also easily detected in the tissue (Figs. I-K). Some branching patterns were nearly indistinguishable from those observed with high resolution in primary serotonergic neuron cultures (Hingorani et al., 2022). Overall, the transgenic model achieved reliable isolation of single serotonergic fibers in a number of brain regions.

**Figure 1.**
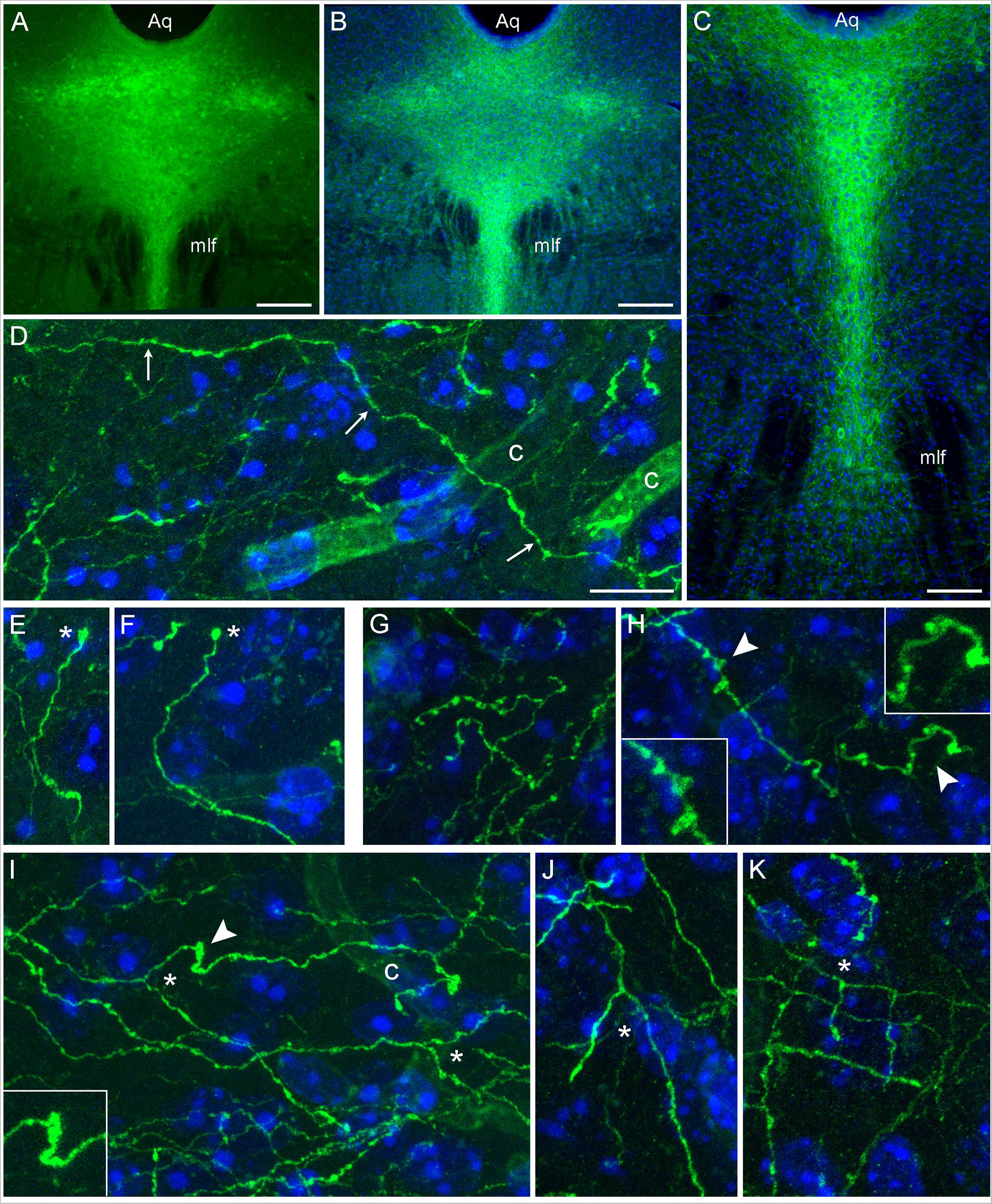
Serotonergic somata and fibers in mice with serotonergic neuron-specific EGFP expression (*Tph2*-*iCreER*; *ROSA*^mT/mG^). These mice were allowed to survive for 10 days after the tamoxifen treatment and were 5 months old at the time of the tissue collection. (**A**) An epifluorescence image of unstained EGFP-positive somata (green) in the dorsal raphe (DR). (**B**, **C**) Epifluorescence images of GFP-positive somata (green) visualized with GFP-immunohistochemistry in the rostral (B) and caudal (C) DR. Cell nuclei are blue (DAPI). (**D**-**K**) Confocal images of GFP-positive fibers (green) visualized with GFP-immunohistochemistry in the superior colliculus (D, E), the inferior colliculus (G, I), and the polymorphic layer of the dentate gyrus of the hippocampus (F, H, J, K). Cell nuclei are blue (DAPI). The images show an isolated fiber (A; arrows), growth cone-like dilated structures (E, F; asterisks), varicosity-like structures (G, H), and branching (I-K; asterisks). The 3D-connectivity of the branching points was verified in the corresponding confocal z-stacks. Some elements (arrowheads) are shown enlarged in insets. Aq, aqueduct; c, capillary; mlf, medial longitudinal fasciculus. Scale bars = 200 µm in (A, B), 100 µm in (C), and 10 µm in (D) (applies to (D-K)).

### 3.2. The Isolation of Single Fibers with Brainbow Labeling

In high-throughput applications, reliable isolation of single fibers should be combined with the visualization of many fibers in the same volume. Simultaneous visualization of many fibers is also important in studies of interactions among different fibers, including the extent of their spatial intermingling (*e.g.*, a small tissue volume may contain fiber segments from many individual fibers or may be dominated by segments from the same few, tortuous fibers). A technical solution is offered by the Brainbow 3.2 toolbox (Cai et al., 2013) which is uniquely well suited for analyses of serotonergic fibers. To our knowledge, it has not been used in this system; an early version of Brainbow (Brainbow 1.0) has been used to visualize serotonergic somata in raphe nuclei (Weber et al., 2009), but it does not support neurite labeling.

In the initial approach, we induced the *Tph2*-dependent Cre-recombination in mice carrying the Brainbow 3.2 transgene (genotyped offspring of *Tph2*-*iCreER* and *Brainbow* crosses) and used immunohistochemistry to visualize the expression of the Brainbow fluorophores in the rostral raphe nuclei. We observed no Brainbow signal, which was likely caused by a low or negligible expression of *Thy1* in the raphe region of this transgenic line, perhaps due to a position effect (Dr. Dawen Cai, personal communication). The *Thy1* gene supports strong transgene expression in many, but not all, neuron types (Cai et al., 2013), and another study has reported a weak or undetectable brainstem expression of a different transgene driven by the *Thy1* promoter (Dana et al., 2018).

To directly test this prediction, we performed an immunohistochemical staining for PhiYFP, a reporter of the transcriptional activity of the Brainbow transgene region before Cre-recombination (Cai et al., 2013). Strongly PhiYFP-positive cells were found in the neocortex and the hippocampus, but no immunoreactivity was detected in the rostral raphe nuclei and adjacent midbrain structures (Fig. 2). This finding can inform other studies attempting Brainbow 3.2 labeling in this neuroanatomical region.

**Figure 2.**
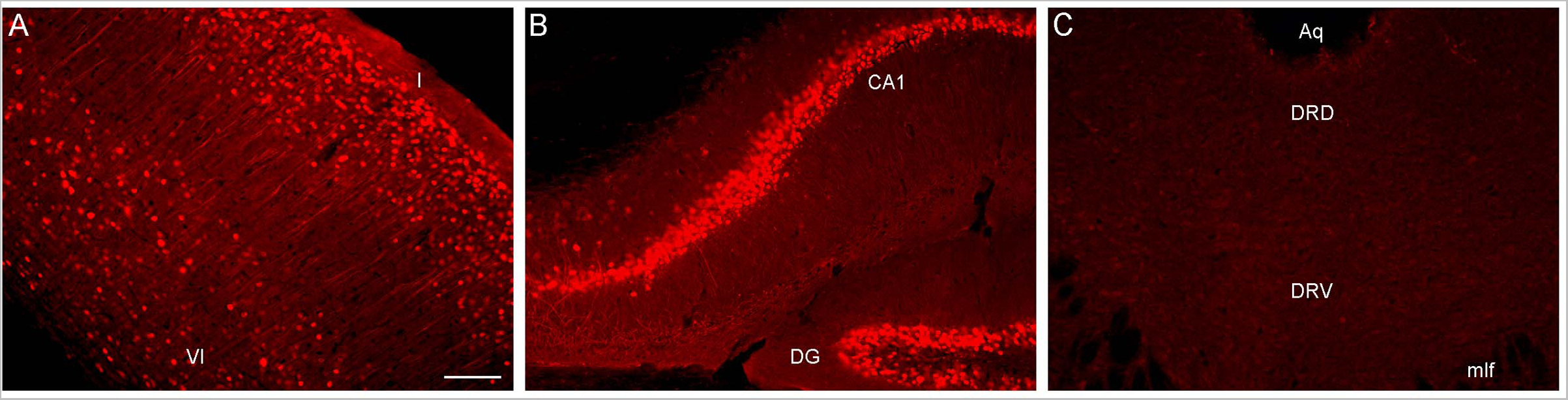
Epifluorescence images of PhiYFP-immunoreactivity in (**A**) the somatosensory cortex, (**B**) the hippocampus, and (**C**) the dorsal raphe (DR) of a mouse with the *Brainbow 3.2* transgene. The PhiYFP-signal is absent from the DR. I and VI, cortical layers I and VI; Aq, aqueduct; CA1, field CA1 of the hippocampus; DG, dentate gyrus; DRD, dorsal DR; DRV, ventral DR; mlf, medial longitudinal fasciculus. Scale bar = 100 µm.

We next attempted an alternative approach, by injecting Brainbow AAVs (Cai et al., 2013) directly into the dorsal raphe of *Tph2*-*iCreER* mice (with the induction of the *Tph2*-dependent Cre-recombination one week later). An immunohistochemical staining for three Brainbow fluorophores (EGFP, mTFP, and TagRFP) revealed labeled somata that were restricted to the raphe complex and showed varying fluorophore-intensity combinations across individual cells, as expected (Fig. 3). Importantly, strong and dense labeling of serotonergic fibers was found in the entire brain of some mice, including the diencephalon and telencephalon (Fig. 4). Generally, the Brainbow-labeling of fibers showed a considerable variability across animal cases, likely because of the uncontrolled differences in the stereotaxic injections, the efficiency of the AAV-transfection, the efficiency of the tamoxifen-induced Cre-recombination, and other experimental and physiological factors. Our best case was an adult male with a post-induction (transport) time of 3 months. The reliability of this approach may be further improved; at this time, we recommend maximizing the probability of a successful outcome by using batches of 5-10 animals. It should be noted that a single successfully labeled brain can provide an enormous amount of information about individual serotonergic fibers in all brain regions, including their spatial interplay.

**Figure 3.**
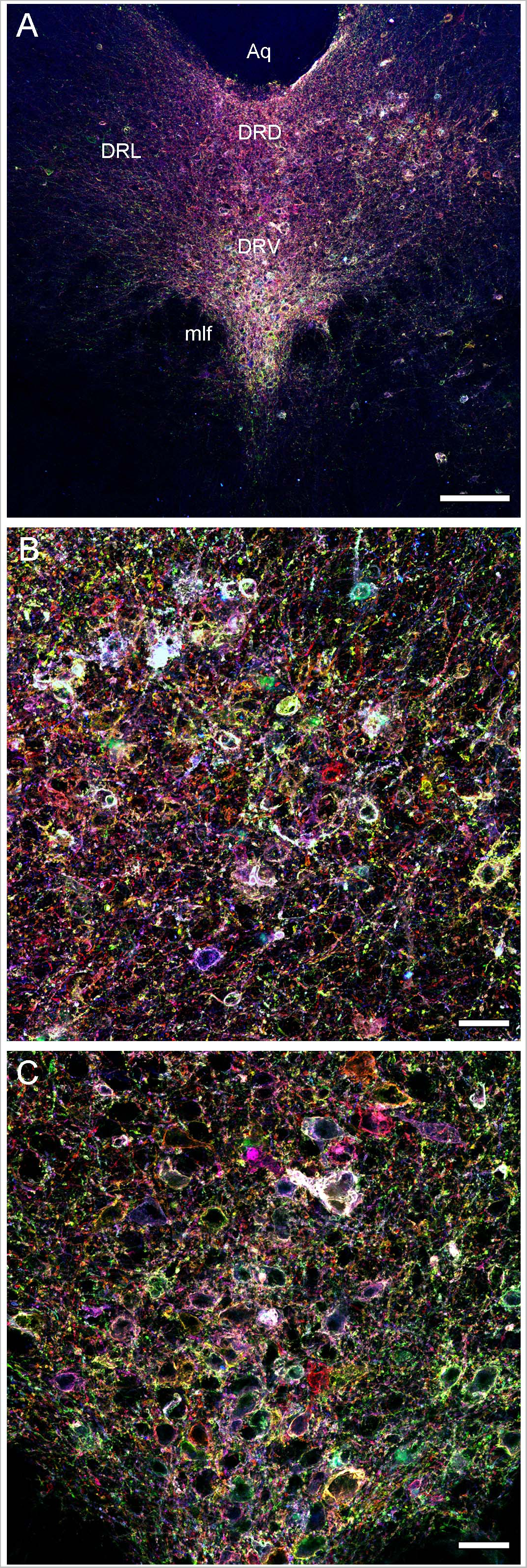
Confocal images of the immunoreactivity of the three Brainbow fluorophores (with the red, green, and blue channels merged) in the dorsal raphe (DR) of a *Tph2*-*iCreER* mouse (male) that received an intracranial injection of the *Brainbow*-AAVs into the DR. This mouse was allowed to survive for around 3 months after the tamoxifen treatment and was around one year old at the time of the tissue collection. (**A**) A low-power image of the DR. (**B**) A high-power image of the dorsal DR. (**C**) A high-power image of the ventral DR. Aq, aqueduct; DRD, dorsal DR; DRL, lateral DR; DRV, ventral DR; mlf, medial longitudinal fasciculus. Scale bars = 150 µm (A), 30 µm (B, C).

**Figure 4.**
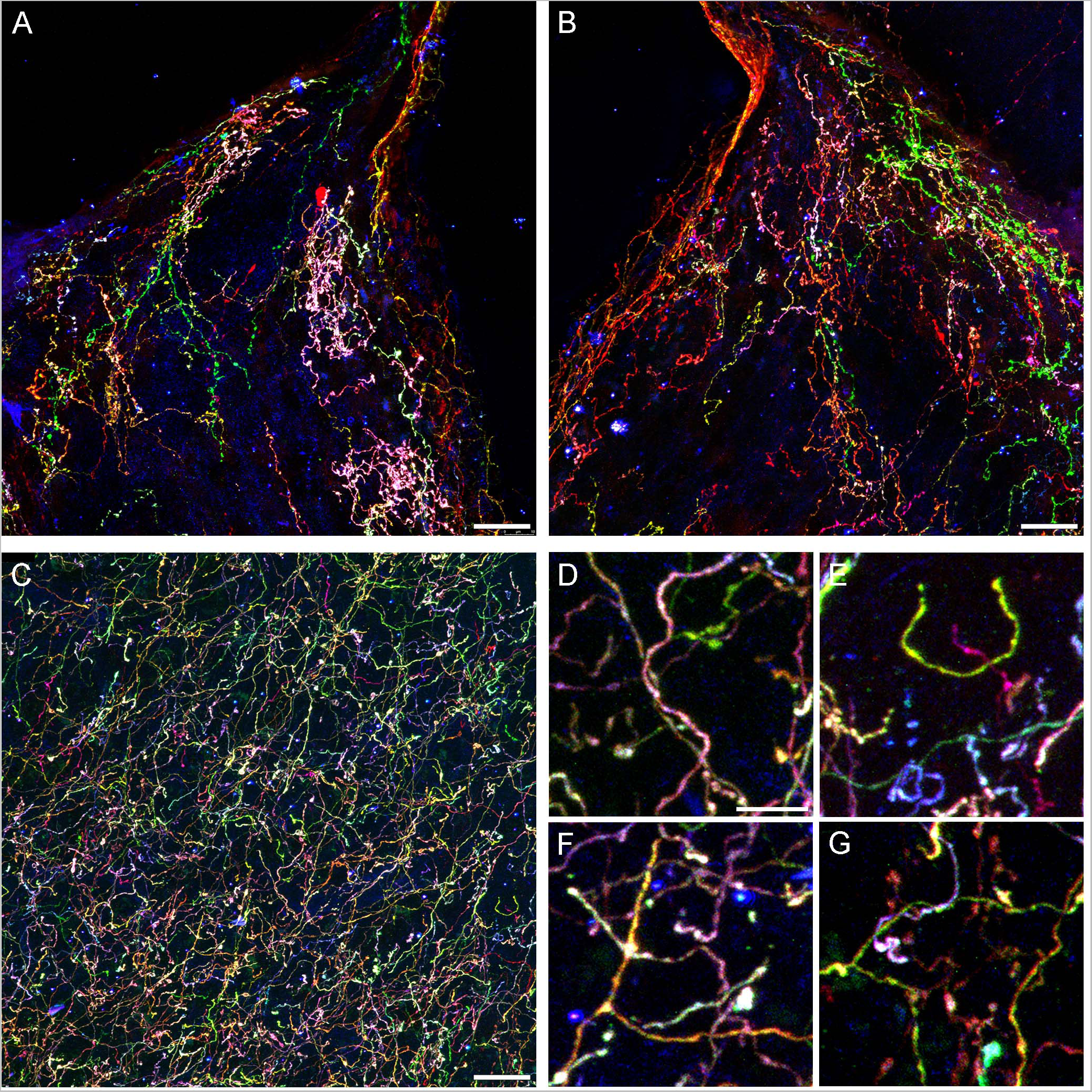
Confocal images of fibers immunoreactive for the three Brainbow fluorophores (with the red, green, and blue channels merged) in the diencephalon and telencephalon of a *Tph2*-*iCreER* mouse (male) that received an intracranial injection of *Brainbow*-AAVs in the dorsal raphe (DR). This mouse was allowed to survive for around 3 months after the tamoxifen treatment and was around one year old at the time of the tissue collection. (**A**, **B**) The left and right habenulas in the same section. In this geometrically constrained space, individual fibers appeared to produce local high-density islands (pink and green in the image). (**C**) The basolateral amygdala. (**D-G**) Four enlarged parts of (C) that show highly tortuous but readily separable trajectories (D, E) and unambiguous branching points (F, G). Scale bars = 20 µm (A-C) and 5 µm (D-G).

Some individual fibers appeared to claim considerable territories in the habenula, a nucleus unusual in its highly constrained geometry (Fig. 4A, B). It is possible that fibers can become trapped in this region, producing highly tortuous trajectories as they interact with the physical borders. In contrast, fibers appeared to be well intermixed in the basolateral amygdala (Fig. 4C), a region with one of the highest densities of serotonergic fibers in the entire brain (Awasthi et al., 2021). Despite this high density, the Brainbow labeling revealed clearly identifiable individual paths (Fig. 4D), including full turns (Fig. 4E), as well as unambiguous branching events (Fig. 4F), some followed by an immediate intermingling of the branches with fibers of different identities (Fig. 4G).

### 3.3. The Quantification of Fiber Trajectories: Theoretical Considerations

This study focuses on one essential characteristic of serotonergic fibers, their trajectories in the three-dimensional space. Considering the individual uniqueness of these trajectories, their rigorous quantification requires some theoretical assumptions.

We started with a top-down approach, by producing a series of computer simulations of serotonergic fibers modeled as step-wise random walks based on the von Mises-Fisher (vMF) directional probability distribution (Fig. 5). This distribution is parametrized by the unitless concentration parameter (κ) that determines the stiffness of the fiber (low κ values make the fiber more flexible, high κ values make it stiffer).

**Figure 5.**
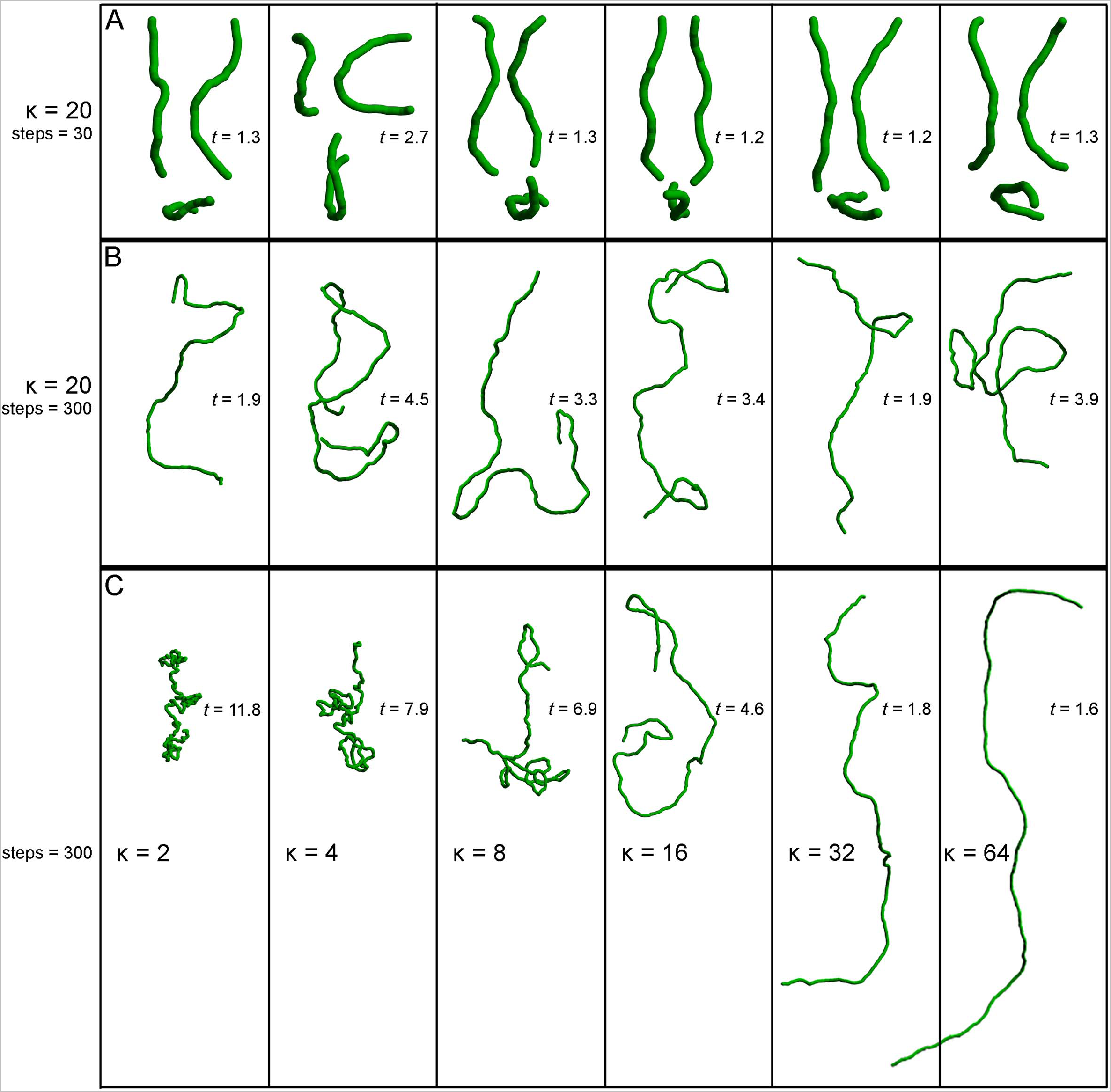
Simulated fibers produced by a step-wise random walk, in which the direction of each step was drawn from the von Mises-Fisher probability distribution with the concentration parameter κ. The diameter of the fibers is 1 µm and each step is 1.5 µm in length. The tortuosity index (*t*) of each fiber, calculated as the fiber length divided by the Euclidean distance between the fiber ends (Jin et al., 2016; Pratelli et al., 2017), is also shown. (**A**) Six realizations of a walk with κ = 20 and 30 steps (the fiber length of 45 µm), approximately corresponding to the mean segment length in a 40 µm-thick section (Janušonis et al., 2019). Each fiber is shown in three non-perpendicular orientations that demonstrate caveats of two-dimensional interpretations (the same fiber may appear very different, depending on its orientation with respect to the imaging plane). (**B**) Six realizations of a walk with κ = 20 and 300 steps (the fiber length of 450 µm). The tortuosity index can be suboptimal in that it can strongly vary in fibers produced by the same stochastic process. (**C**) Single realizations of walks with six κ values (2 – 64) and 300 steps (the fiber length of 450 µm).

The selected model was based on our previous theoretical investigations (Janušonis and Detering, 2019; Janušonis et al., 2019). The shown simulations are more physically realistic in that they include an accurate scaling of the fiber diameter (set to 1 µm) with respect to its length and also use a physically-informed step (set to 1.5 µm). The simulations show the typical appearance of fiber segments in 40 µm-thick sections (Fig. 5A), computational extrapolations of fibers over longer distances, with the corresponding tortuosity indices (Fig. 5B), and the dependence of the fiber appearance on the concentration parameter, again with the corresponding tortuosity indices (Fig. 5C). The model performs well in capturing the geometry and inherent variability of single serotonergic fibers. In addition, the simulations demonstrate the limitations of the tortuosity index which can vary strongly in different realizations of the same process (Fig. 5B).

Since fibers are assumed to advance in steps that maintain the same length (but not the same direction), the step length should be ideally optimized with an unbiased and physically-informed procedure. It is easy to see at the qualitative level that an excessively small step can make a fiber appear stiffer that it actually is, and that an excessively large step may lead to unstable computational estimates, because the step can skip over important direction changes (Fig. 6A). This problem can become more severe with noise, especially if the step and the noise signal are comparable in magnitude (Fig. 6B). One solution is to always indicate the step at which the concentration parameter was estimated (*e.g.*, κ(2 µm) = 30), but such arbitrary decisions can complicate quantitative comparisons among different studies.

**Figure 6.**
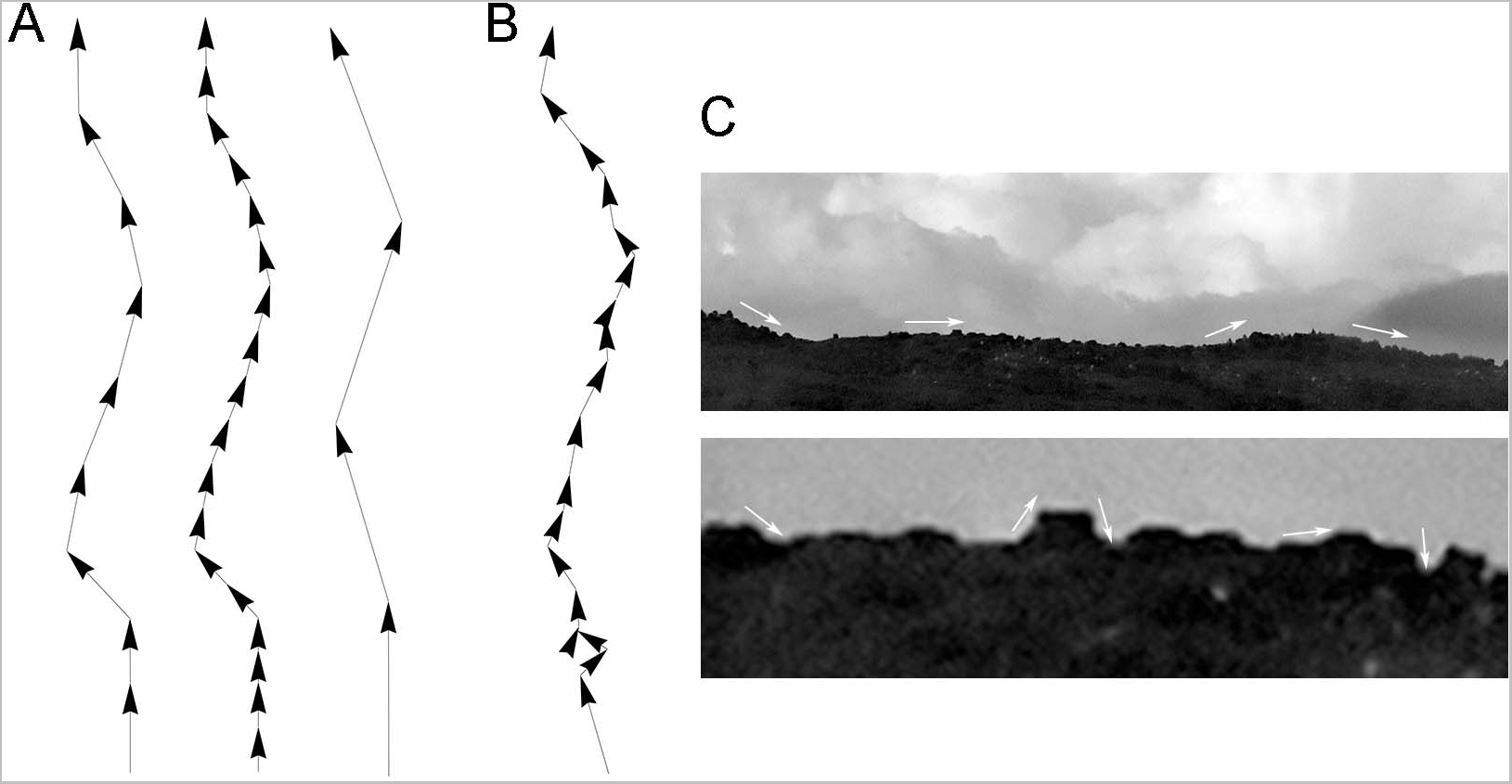
Qualitative effects of the sampling step along a fiber trajectory. (**A**) Fiber trajectories can be described with a natural step that reflects the physical structure of the fibers (**left**). With no noise, *smaller* steps (oversampling) will make a fiber appear more rigid and produce higher concentration parameter (κ) values (**middle**). In contrast, *larger* steps (undersampling) can make a fiber appear more flexible and produce lower κ values; but it can also result in computationally unstable directional distributions (**right**). (**B**) With noise, unnaturally small steps can also become unstable because their order of magnitude may become comparable to that of the noise. (**B**) This phenomenon is demonstrated with the contour of mountains, where an unnaturally small sampling step may capture the contour of relatively small, irrelevant objects and grossly reduce the estimate of κ (*i.e.*, the contour may appear less rigid than it actually is).

First, we simulated a long fiber, with a known value of the concentration parameter (κ = 20), and computed the *estimates* of this parameter with the original step (set to one unit), as well as with fractional steps and integer-multiples of the original step (Fig. 7A). As expected, the real parameter value was recovered at step = 1. However, the parameter was grossly overestimated with the smaller steps and slight underestimated with the larger steps. It should be noted that we did not include extremely large steps (*e.g.*, jumping from the beginning of the fiber to its mid-point), which would make the estimates unstable. Importantly, the plot produced an “inflection” point, corresponding to the correct step length.

**Figure 7.**
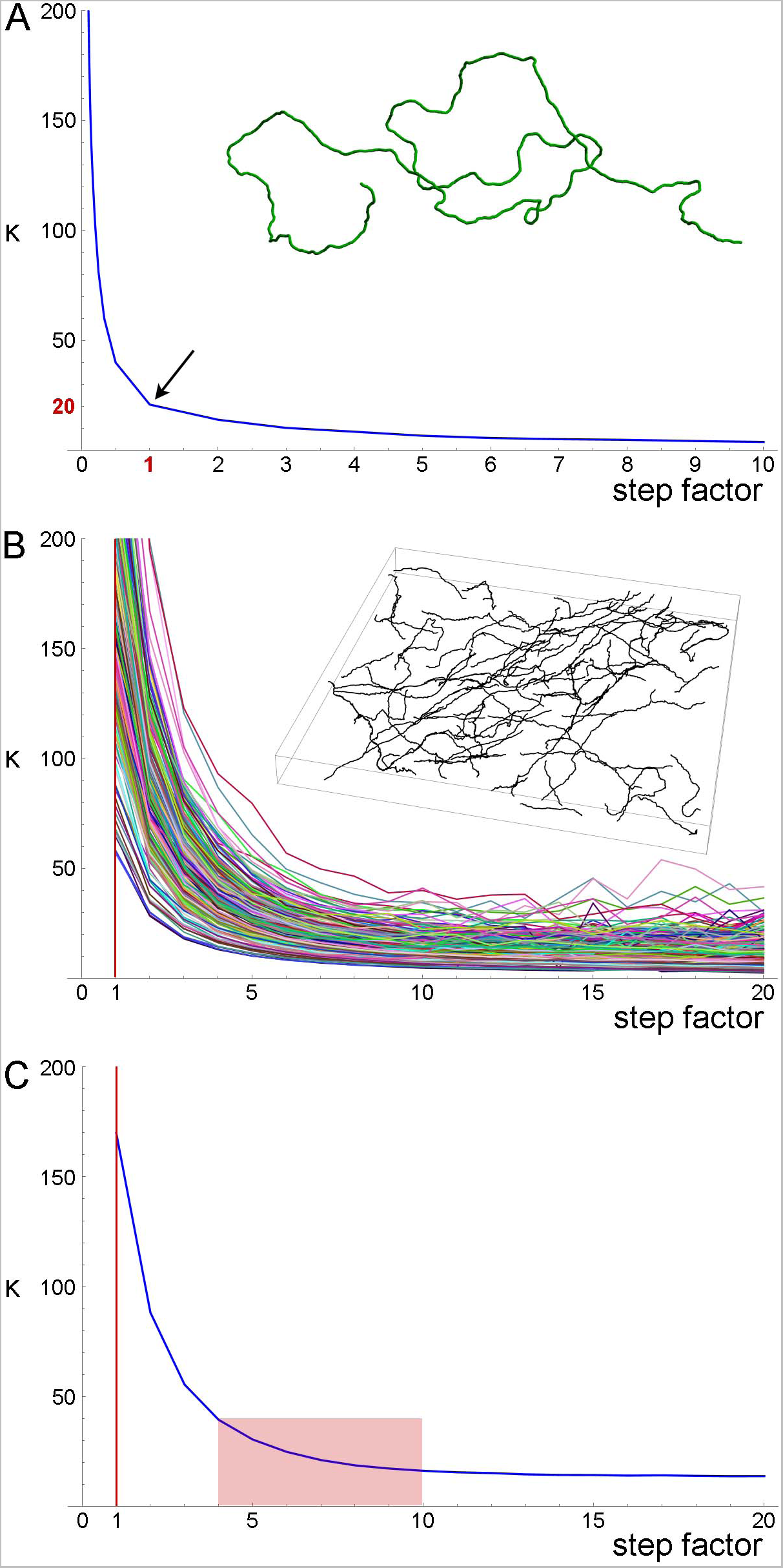
A quantitative method to obtain an optimal step for the sampling of fiber trajectories. (**A**) The *estimated* κ values of a simulated fiber (shown in the inset) with the *actual* κ = 20. The fiber length is 1000 steps (one length-unit each). The smaller steps were produced by dividing each original step into 2, 3, … 9, and 10 steps (producing steps of 0.1 – 0.5 units), and the larger steps were produced by multiplying the original step by factors of 2 – 10 (producing steps of 2 – 10 units). Even if the original step were unknown, it could be recovered at the sharp “inflection” of the plot. (**B**) The estimated κ values of 213 serotonergic fibers in the mouse somatosensory cortex that have been visualized with 5-HT-immunohistochemistry (using a published protocol (Janušonis, 2018)), imaged with confocal microscopy, and manually traced through three-dimensional z-stacks with Simple Neurite Tracer, an ImageJ plugin (the inset shows traced fibers in one z-stack). The plugin automatically selects a step of approximately 0.15 µm (Janušonis and Detering, 2019), which is considerably smaller than a typical fiber diameter. A gradual increase in the step length shows that the original step strongly overestimates κ. (**C**) The mean estimated κ values of the fiber set in (C), which reveals an “inflection region” at the factor values of around 3 – 10. Considering the diameter of a typical fiber (around 1 µm), the factor of 10 (corresponding to a step of approximately 1.5 µm) is a well-balanced choice.

Second, we attempted the same approach in a set of actual serotonergic fibers that have been manually traced by several trained individuals blind to the model (Fig. 7B). In this case, the “correct” step length was not known but could be recovered in a similar “inflection region.” The plot of the population means (Fig. 7C) put it at around 0.6 – 1.5 µm, which is in register with the typical diameter of serotonergic fibers. With additional physical and computational considerations, we set the step at 1.5 µm in the following analyses.

### 3.4. High-precision Tracing of Single Fibers with a Novel Algorithm

After a fiber has been isolated with transgenic methods and imaged with high resolution in the three spatial dimensions, it has to be converted to an array of XYZ-coordinates (*i.e.*, an *N* × 3 matrix, where *N* is the number of points). These coordinates (more precisely, the resultant step-vectors) are sufficient to produce a numerical estimate of the concentration parameter. This estimate becomes more accurate with larger step numbers, with one important caveat: all steps should reliably belong to the same fiber (*i.e.*, the trace cannot accidentally jump to another, adjacent fiber). Identity errors cannot be easily mitigated with larger samples; a single long, accurate trace can produce a more reliable estimate than a set of unreliable traces.

The conversion of visualized serotonergic fibers to trajectory arrays is a non-trivial problem. Manual tracing is highly unreliable because human tracers typically have to move up and down z-stacks (a significant perceptual and memory challenge in noisy images), have to make subjective decisions in fibers whose varicosities alternate with signal interruptions, and cannot maintain a steady level of performance due to fatigue. General-purpose systems, such as Imaris (Bitplane) have no built-in knowledge of the physical properties of serotonergic fibers, which (in our hands) results in frequent incorrect decisions. Deep learning-based segmentation systems can separate serotonergic fiber segments from the background, also in whole-brain images, but they currently cannot reliably trace the *individual*, uninterrupted trajectories of fibers over useful distances (Friedmann et al., 2020).

We developed an algorithm that overcomes some of these difficulties. It prioritizes accuracy over volume and speed and is designed for high-resolution, three-dimensional confocal images of serotonergic fibers labeled with a single fluorophore. In particular, it is highly appropriate for EGFP-labeled serotonergic fibers in the described transgenic model. In the near future, it can be extended to Brainbow-labeled fibers (with an incorporation of the color dimension). The key components of this algorithm are shown in Figure 8. We next describe it in detail.

**Figure 8.**
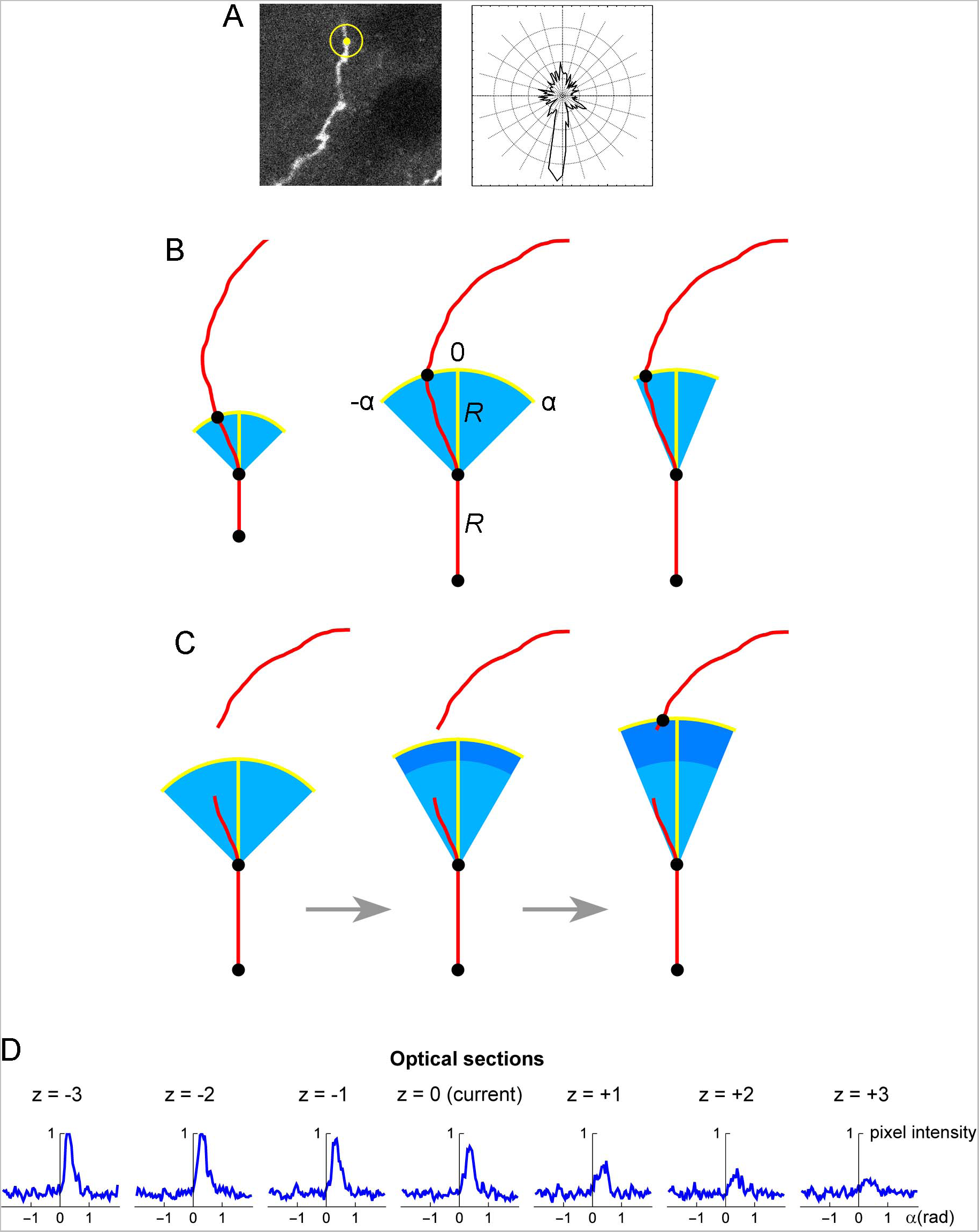
(**A**) A seed point (yellow) and the initial search circle around it (with a radius of around 2 µm; **left**), with a corresponding polar plot of the brightness values on the circle (where larger radii correspond to stronger brightness; **right**). The polar plot indicates the position of the second point, which initiates the automated tracing procedure. (**B**) At each current fiber point (an XY-coordinate in a specific optical section), the “flashlight” tracing algorithm scans the intensity of the pixel values in an arc that is centered at this point, with a radius *R* and the subtended angle [-*α*, *α*]. The middle point of the arc (0) corresponds to the “expected” direction of the fiber (marked with the yellow segment), which is based on the current point and the point before it. The values of *R* and α can be adjusted to reflect the general tortuosity of the fiber. For clarity, the first segment is shown straight and the value of *R* is greatly exaggerated (it is typically on the order of the fiber width). A specified number of optical sections above and below the current optical sections are also scanned in the same way. (**C**) If the algorithm fails to detect any intensity peaks in the given subset of optical sections, the step can be automatically extended (the dark blue region). Since this extension increases the risk of false-positives (*e.g.*, it may detect a segment of another fiber in the vicinity), the search angle can be automatically decreased. The extended *R* is an integer multiple of the original *R* (0.5*R*, 1*R*, 1.5*R*, 2*R*, …); in the figure, it is shown smaller for compactness. (**D**) An example of a decision made by the tracing algorithm in one step (taken from the tracing sequence of an actual EGFP-labeled fiber). The center plot shows the pixel intensity values in the current optical section (*z* = 0) in the arc centered at the current XY-position with *R* = 2 µm and α = 0.6π radians (the current direction is given by *α* = 0). The plots to the left and right of the middle plot show the corresponding pixel values at the same current XY-position and direction in the three optical sections below and above the current optical section (typically, more adjacent sections are used). In this example, the trace turns slightly to the right and three optical sections down. The trace then continues iteratively with the new XY-position, optical section, and direction.

The *z*-stack of one fluorophore channel is imported as a series of high-resolution, grayscale TIFF images (with the XY-scaling of around 60 nm per pixel and the distance between two adjacent optical sections of around 300 nm). The images are auto-contrasted (with the brightness range of 0-1) and Gaussian-blurred using a one-pixel radius. The initial seed point is selected on a fiber in a specified (“current”) optical section, manually or automatically, and the second point is automatically detected as the brightest point on a circle whose center is at the seed point and the radius is set at a fixed value (*R*). The second point becomes the “current point,” and the “current direction” is determined by the vector connecting the two points. This initiates the tracing sequence (Fig. 8A).

In the current optical section, the brightness of pixels is examined on an arc that is centered at the current point. The radius (tracing step) and angle of the arc are given by *R* and a fixed interval [-*α*, *α*], respectively, where the angle of 0° corresponds to the current direction (Fig. 8B). The number of increments (2*n*) from −*α* to *α* should provide a sufficiently dense coverage of the arc; in particular, the arc increment (*α*/*n*) should be less than one-half of the expected fiber width. By sampling the arc, brightness peaks within a fixed brightness interval [*B_min_*, *B_max_*] are detected, but the peaks whose width (spatial extent) at *B_min_* falls below a set threshold (*W_min_*) are not kept. The latter procedure, along with the described Gaussian blur, efficiently eliminates pixels whose strong brightness is noise-related and does not extend to adjacent pixels. In the remaining peaks, the peak *closest to the current direction* is selected (if there is more than one), and its point with the brightest value is recorded as the *next candidate point*.

In addition to the current optical section, this procedure is repeated in *N* sections above and *N* sections below the current optical section (if they exist), using the same arc. Among the next candidate points across the entire subset of optical sections, the *brightest point is selected*. It becomes the next current point of the fiber. The next current direction is determined by the vector connecting the XY-coordinates of the two most recent points, and the procedure continues iteratively. It is automatically terminated when brightness peaks are no longer detected or when an edge of the stack is reached. It can also be terminated manually, after a fixed number of steps.

We note that the algorithm is “anisotropic” in that within each optical section it prefers a bright value that is closest to the current direction, but among the optical sections it prefers the brightest value in the set (which may not be closest to the current direction in all sections). This choice was based on tests in actual z-stacks and it reflects an inherent property of confocal imaging: confocal images are sets of 2D-images, not true 3D-images with an isotropic resolution.

As described, the algorithm would not be able to handle transient interruptions in signal brightness that accidentally fall on the search arc. To prevent this, the algorithm can automatically extend its search radius to multiples of *R* (0.5*R*, 1*R*, 1.5*R*, 2*R*, …), if no brightness peaks are detected across the optical section subset (Fig. 8C). The set includes one radius that is smaller than the original radius. However, the increasing extensions (numbered *i* = 1, 2, 3, 4, …, *i_max_*, where *i* = 2 corresponds to the original *R*) increase the risk that the trace will jump to another fiber located close to the fiber that is being traced. Therefore, these extensions can be optionally accompanied by an automatically narrowing search angle, given by *α*(*i*) = *α_o_e*^−*k*(*i*−2)^, where *α_0_* is the original *α* and *k* is a constant (the “narrowing strength”). The maximal number of extensions should be restricted (*i_max_*) to a small value, especially in dense areas. If the next point is detected with the extensions, in the next step the extended *R* and the narrowed *α* revert to their original values.

Since the arc is set in a 2D-plane, the actual distances between adjacent points in the 3D-trace may slightly differ from the original *R*. Extended *R* values also contribute to these deviations. However, final traces can be automatically fine-tuned with a 3D-step of a fixed length, by simply sliding along the original trace and re-marking (“nudging”) all points such that they are separated by the same exact Euclidean distance. Since the original steps are small, the recalculated trajectory is typically visually indistinguishable from the original one. However, the constant step improves the reliability of concentration parameter estimates.

In some instances, the algorithm may fail to correctly process a step. It can then be manually forced to bridge the gap, with an automatic continuation. With an optimal parameter set, such instances should be rare. In some cases, these adjustments are necessary to make the trace follow a specific branch, when two options are available. The question of which branch is the “correct” one poses interesting questions in stochastic analyses.

Figure 8D shows an example of the automatic detection of the next trace point in an actual EGFP-labeled serotonergic fiber. The typical parameter values are given in Table 1. The algorithm is currently implemented in Wolfram Mathematica, a leading computer language in functional programming, but it can be easily moved into other languages such as Python or Java.

**Table 1.** Typical values of the tracing algorithm. The values were obtained in stacks of grayscale images of 3144 × 3144 pixels, with the XY-scaling of 60 nm/pixel and the Z-step of 300 nm. In some ranges, more frequent values are shown in bold.

| Parameter | Meaning | Approximate value |
| --- | --- | --- |
| $R$ | default search radius (tracing step) | 1.5-2.0 $\mu\text{m}$ |
| $\alpha = \alpha_0$ | default search angle (one side) | $\sim \pi/2$ ($90^\circ$) |
| $n$ | number of angle increments (one side) | $\sim 50$ |
| $k$ | angle attenuation in extensions | 0.0-0.1 |
| $i_{\max}$ | maximal number of extensions | **0-4** |
| $B_{\min}$ | minimal brightness | $\sim 50\%$ |
| $B_{\max}$ | maximal brightness | $\sim 100\%$ |
| $W_{\min}$ | minimal peak width | 0.1-0.2 $\mu\text{m}$ |
| $W_{\max}$ | maximal peak width | 1.0-**3.0** $\mu\text{m}$ |
| $N$ | number of neighboring sections (one side) | $R/(\text{z-step})$ |

### 3.5. The Quantification of Fiber Trajectories: An Experimental Example

After the image of a fiber has been converted into an array of XYZ-coordinates, with a constant step in the 3D-space, the trace can be iteratively rotated such that the current step always coincides with the unit-vector (0, 0, 1). The direction of the next step is then recorded as a unit-vector, and the next step becomes the current step (Fig. 9). The length of the *mean* of the collected vector sample (shown in red in Figure 9) is used to calculate an estimate of the vMF concentration parameter (κ), as described in detail in the Materials and Methods.

**Figure 9.**
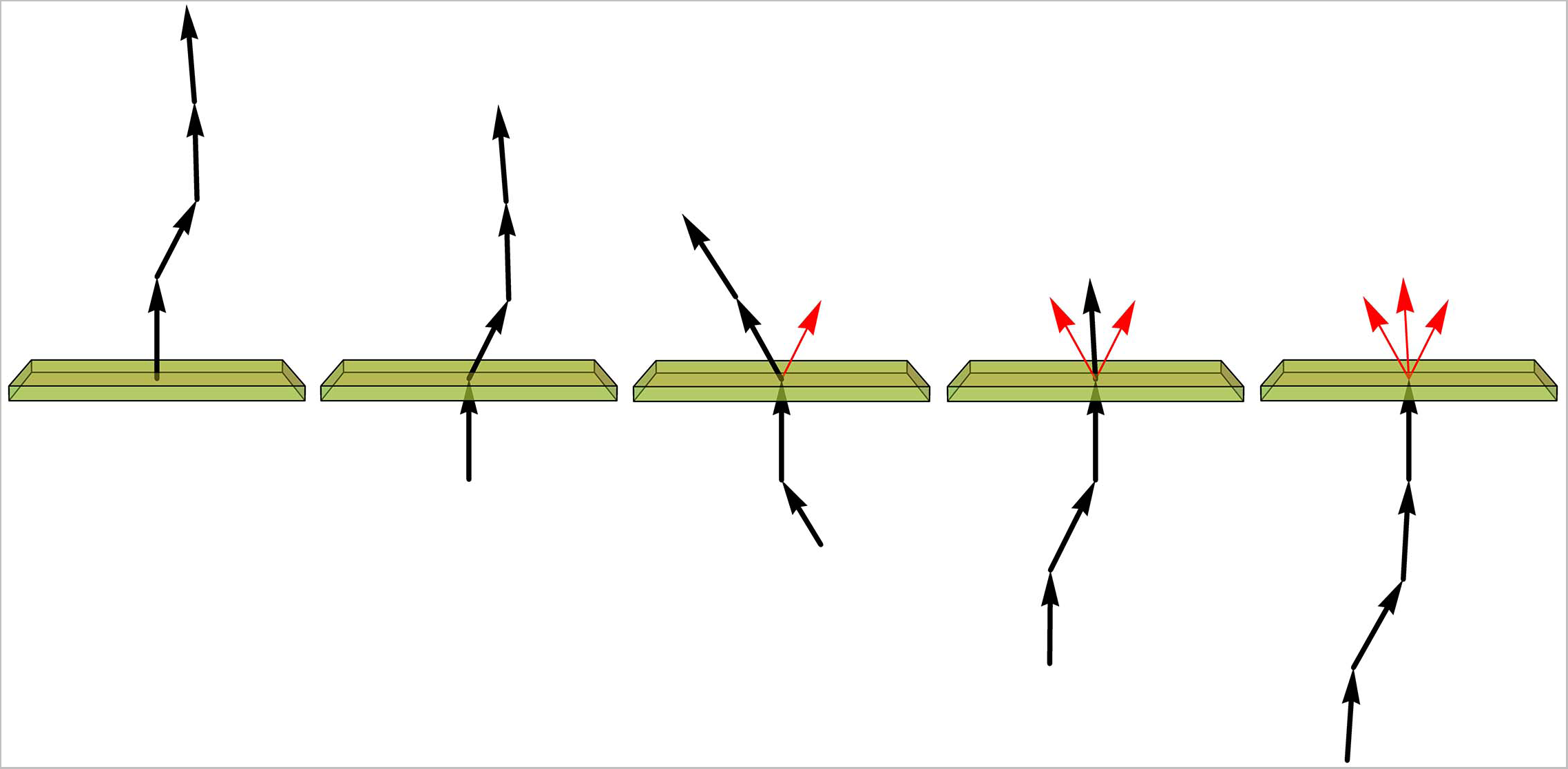
Assuming the fiber is composed of constant-length segments, each of which has a direction drawn from the von Mises-Fisher distribution (with the expected direction of each step given by the direction of the previous segment), the value of the concentration parameter (κ) can be estimated by sequentially rotating each current segment (a unit-vector) to the vertical position (shown just below the plane), recording the direction of the next segment (represented by the unit vector just above the plane), and calculating the length (*L*) of the mean of all direction vectors (shown in red). If *L* > 0.9, κ = 1/(1 − *L*) is a good approximation. In particular, if all direction vectors point in the same direction, *L* approaches 1 and κ approaches infinity, as expected for a perfectly rigid, straight fiber.

We computed the concentration parameters in a sample of EGFP-labeled fibers in the inferior colliculus of mice with the *ROSA*^mT/mG^ transgene, after the *Tph2*-dependent Cre-recombination (Fig. 10). Most of the values were tightly clustered around κ = 20 (for the used step of 1.5 µm), but two of them were around κ = 40. Larger samples are required to achieve a better understanding of the distribution of this parameter, which may reveal differences among fibers belonging to transcriptionally different neurons or located in different brain regions. For example, a recent study has found two functionally distinct types of serotonergic fibers in the hippocampus (Luchetti et al., 2020). Such analyses fall outside the scope of this methodological study, but we are currently pursuing some of these directions.

**Figure 10.**
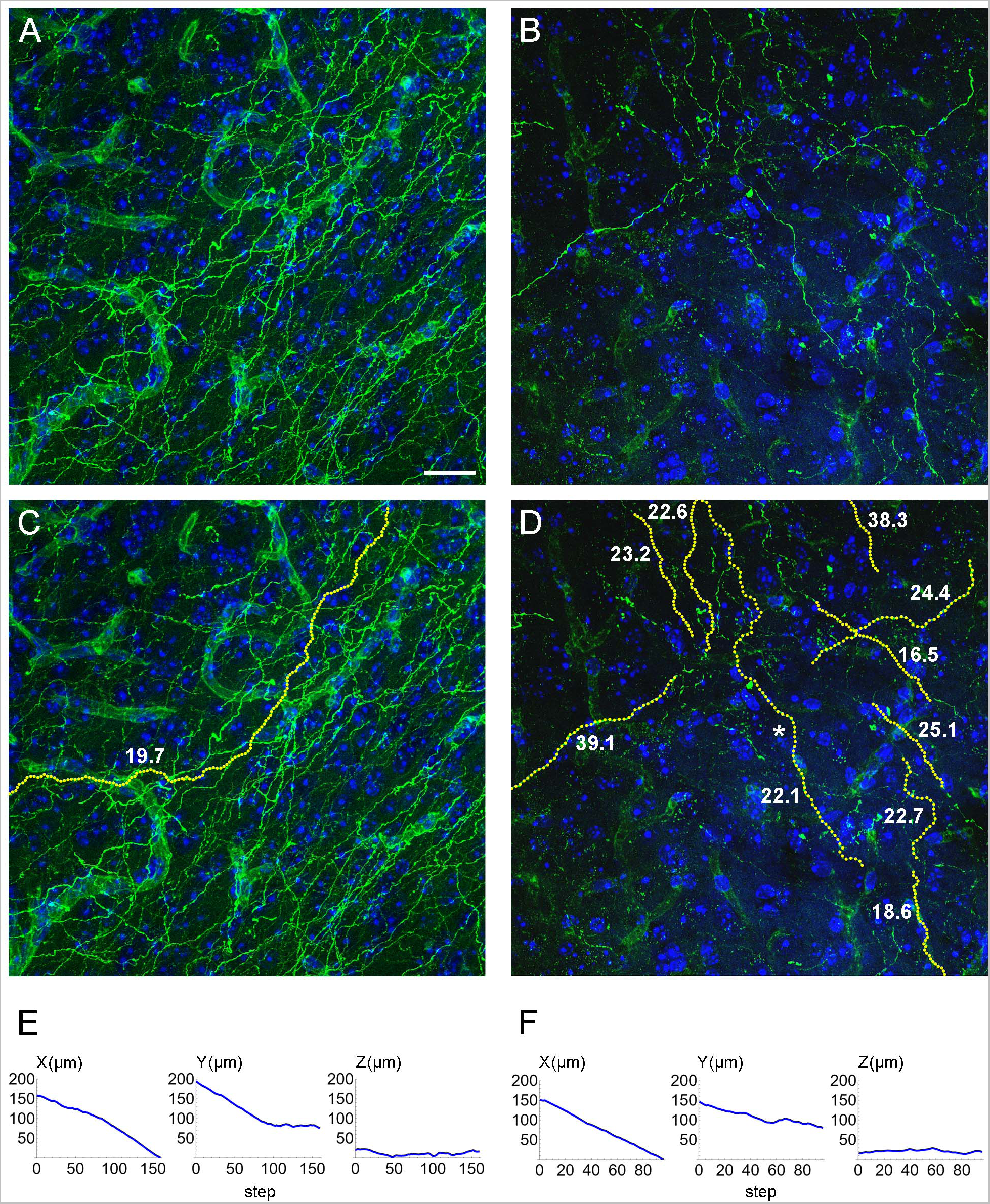
(**A**, **B**) Two confocal images of GFP-immunoreactive fibers in the inferior colliculus of mice with serotonergic neuron-specific EGFP expression (*Tph2*-*iCreER*; *ROSA*^mT/mG^). (**C**) An automatically traced fiber in the z-stack corresponding to (A). The fiber is located in an area with a relatively high fiber density, but the algorithm can still follow its trajectory because fiber intersections can be disambiguated in the 3D-volume. (**D**) A set of automatically traced fibers in a relatively noisy z-stack corresponding to (B). In (C) and (D), each yellow dot marks a detected trajectory point. The tracing quality was verified with visual examination of the traces across all optical sections. The estimated κ values (at the step of 1.5 µm) are shown in bold. (**E**) The X-, Y, and Z-coordinates (as a function of the step number) of the traced fiber in (C). (**F**) The X-, Y, and Z-coordinates (as a function of the step number) of one of the traced fibers in (D) (marked with an asterisk). In (E) and (F), the Z-coordinate is constrained within a range of 40 µm (because of the thickness of the physical sections). Scale bar = 20 µm.

## 4. DISCUSSION

We developed an integrated system for analyses of single serotonergic fibers in brain tissue. It includes two transgenic mouse models for the visualization of individual fibers, a mathematical model of the fiber trajectories, a high-precision algorithm for tracing fibers in 3D-images, and procedures to estimate a single theoretical parameter from complex fiber paths. This system can be used in many experimental setups, including those than include serotonergic perturbations, and can be readily deployed in any mouse brain regions. The study focused on the rostral raphe nuclei, but the caudal raphe nuclei also express *Tph2*, making this system extendable to the lower brainstem (with serotonergic fibers descending to the spinal cord). Since both mouse models are based on the *Tph2*-dependent Cre-recombination, the single-fluorophore model may express EGFP in the gut because the *Tph2* gene is also active in enteric neurons (Gershon, 2009; Neal et al., 2009; Yabut et al., 2019) and the *ROSA*^mT/mG^ construct is transcribed in the gut (Muzumdar et al., 2007). However, the overwhelming amount of peripheral serotonin is synthesized by the gut enterochromaffin cells that express a different gene, *Tph1* (Gershon and Tack, 2007). Because the Brainbow AAVs are injected intracranially, the labeled somata should be restricted to the raphe complex in this model.

It should be noted that a number of other Cre models have been produced for studies of serotonergic neurons. Some of them may not label all serotonergic neurons and some may label additional neuron populations (Cardozo Pinto et al., 2019). They can be used in combination with various Cre reporters (Navabpour et al., 2020) and may work well with the Brainbow AAVs.

At present no software exists to reliably trace single serotonergic fibers over distances required for statistical estimations (50-100 µm and longer). This includes automated image-segmentation systems based on convolutional neural networks (CNNs) (Friedmann et al., 2020). The training of CNNs crucially depends on large volumes of annotated data; using models pretrained on other filamentous structures is not an option because they are likely to have other physical properties. Such large datasets cannot realistically (and reliably) be generated with manual tracing. We note that Brainbow labeling effectively achieves such annotation because it resolves ambiguities in many points where fibers cross at short distances. Training sets can be further augmented with simulated fibers that can be computer-generated at various densities in unlimited quantities. Once the network has acquired knowledge about what serotonergic fibers can and cannot do, it can be used to automatically analyze tissue samples labeled with a *single* fluorophore, including human brain biopsies and post-mortem samples processed with immunohistochemistry.

We note that currently two mathematical models of serotonergic fibers exist. The first model is based on a step-wise walk guided by the vMF-distribution (Janušonis and Detering, 2019) and was used in this study. The second model is based on fractional Brownian motion (FBM) (Janušonis et al., 2020; Janušonis et al., 2023). Both models appear highly promising but are not mathematically compatible. One essential difference, important in practical applications, is that that the vMF-model is purely spatial and has no time term; in contrast, FBM-paths develop in space as a function of continuous time (notably, with no definable “velocity” at any time point). Since the real-time dynamics are typically unavailable in experimental fiber analyses (in most cases, fibers are studied in fixed preparations), the vMF-model is superior in this case. However, the FBM-model promises to eventually capture the “deep” structure of serotonergic fibers, as they evolve in both space and time. Live-imaging recordings of growing serotonergic fibers are needed to advance these studies; efforts have already been made in this direction (Jin et al., 2016; Hingorani et al., 2022). Interestingly, *normal* Brownian motion (a special case of FBM) and the vMF-distribution *are* related (Mardia and Jupp, 2000; Gatto, 2013). It should also be noted that the field of “active Brownian particles” may offer other theoretical tools (Romanczuk et al., 2012). We are actively investigating the applicability of these frameworks in our larger research program.

Directional statistics deals with probability distributions on manifolds (*e.g.*, circles, spheres) rather than familiar Euclidean spaces (*e.g.*, lines, planes) (Mardia and Jupp, 2000). A trivial example is given by probability distributions on the unit circle, where the same point can be referred to as 0 or 360 (in degree units). Therefore, caution should be exercised in applications of standard statistical tests to concentration parameter estimates. Directional methods exist to test unit-vector samples for their consistency with the von Mishes-Fisher distribution (against other probability distributions), to compare the concentration parameter estimates in two and more samples (which is particularly relevant to fiber comparisons), to perform ANOVA- and regression-like tests, and to carry out other procedures analogous to those is classical statistics (Mardia and Jupp, 2000; Pewsey and Garcia-Portugués, 2021). A number of R packages are available for these calculations (Pewsey and Garcia-Portugués, 2021), of which *Directional* (Tsagris et al., 2023) is particularly useful in the present context.

The proposed system can support the rapidly growing interest in single serotonergic neurons and fibers, far beyond their classical (population-based) conceptualizations. Recent studies include such diverse approaches as single-cell RNA-seq (Okaty et al., 2020), high-resolution analyses in primary brainstem cultures (Hingorani et al., 2022), supercomputing-based modeling (Janušonis et al., 2020; Janušonis et al., 2023), potential applications in artificial neuron networks (Lee et al., 2022), and the development of novel, broadly-applicable models in physics, directly inspired by the properties of the serotonergic axons (Vojta et al., 2020; Saito, 2023; Wang et al., 2023). It is conceivable that these new frameworks will produce new fundamental insights into the brain as a self-organizing system.

## 5. Conflict of Interest

The authors declare that the research was conducted in the absence of any commercial or financial relationships that could be construed as a potential conflict of interest.

## 6. Author Contributions

KCM developed all experimental methods and contributed to all theoretical discussions. JHH made major contributions in assisting KCM in all experimental procedures and in the maintenance of transgenic mouse colonies. SJ initiated and supervised the project, performed all computational analyses, and wrote the manuscript.

## 7. Funding

This research was supported by the National Science Foundation (grants #1822517 and # 2112862 to SJ), the National Institute of Mental Health (#MH117488 to SJ), and the California NanoSystems Institute (Challenge-Program Development grants to SJ). We acknowledge the use of the NRI-MCDB Microscopy Facility and the Leica SP8 Resonant Scanning Confocal Microscope (supported by National Science Foundation MRI grant #1625770).

## 8. Acknowledgements

We acknowledge the technical contributions of Kirra Thomas, George Grant, and Marissa Millwater (members of the Janušonis laboratory). This work benefitted from manual fiber tracing performed at the beginning of the project by Kimia Boldaji, Natalie Mohr, Jenna Sanfilippo, and Stephanie Valdez. We thank Dr. Benjamin Lopez (UCSB Neuroscience Research Institute) and Dr. Jennifer Smith (UCSB California NanoSystems Institute) for their technical support in microscopy imaging and genotyping. We also thank Dr. Dawen Cai (University of Michigan) for his advice in the search for solutions to technical challenges in the Brainbow labeling.

## Notes

### Competing Interest Statement

The authors have declared no competing interest.

## REFERENCES

Awasthi, J.R., Tamada, K., Overton, E.T.N., and Takumi, T. (2021). Comprehensive topographical map of the serotonergic fibers in the male mouse brain. J Comp Neurol 529(7), 1391–1429. doi: 10.1002/cne.25027.

Azmitia, E.C., Singh, J.S., and Whitaker-Azmitia, P.M. (2011). Increased serotonin axons (immunoreactive to 5-HT transporter) in postmortem brains from young autism donors. Neuropharmacology 60(7-8), 1347–1354. doi: 10.1016/j.neuropharm.2011.02.002.

Baas, P.W., and Heidemann, S.R. (1986). Microtubule reassembly from nucleating fragments during the regrowth of amputated neurites. J Cell Biol 103(3), 917–927. doi: 10.1083/jcb.103.3.917.

Baizer, J.S. (2001). Serotonergic innervation of the primate claustrum. Brain Res Bull 55(3), 431–434. doi: 10.1016/s0361-9230(01)00535-4.

Bennis, M., Gamrani, H., Geffard, M., Calas, A., and Kah, O. (1990). The distribution of 5-HT immunoreactive systems in the brain of a saurian, the chameleon. J Hirnforsch 31(5), 563–574.

Bhat, S.K., and Ganesh, C.B. (2023). Organization of serotonergic system in *Sphaerotheca breviceps* (Dicroglossidae) tadpole brain. Cell Tissue Res 391(1), 67–86. doi: 10.1007/s00441-022-03709-7.

Cai, D., Cohen, K.B., Luo, T., Lichtman, J.W., and Sanes, J.R. (2013). Improved tools for the Brainbow toolbox. Nat Methods 10(6), 540–547.

Cardozo Pinto, D.F., Yang, H., Pollak Dorocic, I., de Jong, J.W., Han, V.J., Peck, J.R., et al. (2019). Characterization of transgenic mouse models targeting neuromodulatory systems reveals organizational principles of the dorsal raphe. Nat Commun 10(1), 4633. doi: 10.1038/s41467-019-12392-2.

Carrera, I., Molist, P., Anadon, R., and Rodriguez-Moldes, I. (2008). Development of the serotoninergic system in the central nervous system of a shark, the lesser spotted dogfish *Scyliorhinus canicula*. J Comp Neurol 511(6), 804–831. doi: 10.1002/cne.21857.

Challet, E., Miceli, D., Pierre, J., Repérant, J., Masicotte, G., Herbin, M., et al. (1996). Distribution of serotonin-immunoreactivity in the brain of the pigeon (*Columba livia*). Anat Embryol (Berl) 193(3), 209–227. doi: 10.1007/bf00198325.

Challet, E., Pierre, J., Repérant, J., Ward, R., and Miceli, D. (1991). The serotoninergic system of the brain of the viper, Vipera aspis. An immunohistochemical study. J Chem Neuroanat 4(4), 233–238. doi: 10.1016/0891-0618(91)90015-5.

Cooke, P., Janowitz, H., and Dougherty, S.E. (2022). Neuronal redevelopment and the regeneration of neuromodulatory axons in the adult mammalian central nervous system. Front Cell Neurosci 16, 872501. doi: 10.3389/fncel.2022.872501.

Dana, H., Novak, O., Guardado-Montesino, M., Fransen, J.W., Hu, A., Borghuis, B.G., et al. (2018). *Thy1* transgenic mice expressing the red fluorescent calcium indicator jRGECO1a for neuronal population imaging *in vivo*. PLoS One 13(10), e0205444. doi: 10.1371/journal.pone.0205444.

Daubert, E.A., Heffron, D.S., Mandell, J.W., and Condron, B.G. (2010). Serotonergic dystrophy induced by excess serotonin. Mol Cell Neurosci 44(3), 297–306. doi: 10.1016/j.mcn.2010.04.001.

Daws, R.E., Timmermann, C., Giribaldi, B., Sexton, J.D., Wall, M.B., Erritzoe, D., et al. (2022). Increased global integration in the brain after psilocybin therapy for depression. Nat Med 28(4), 844–851. doi: 10.1038/s41591-022-01744-z.

Dean, J.G., Liu, T., Huff, S., Sheler, B., Barker, S.A., Strassman, R.J., et al. (2019). Biosynthesis and extracellular concentrations of *N,N*-dimethyltryptamine (DMT) in mammalian brain. Sci Rep 9(1), 9333. doi: 10.1038/s41598-019-45812-w.

Economo, M.N., Clack, N.G., Lavis, L.D., Gerfen, C.R., Svoboda, K., Myers, E.W., et al. (2016). A platform for brain-wide imaging and reconstruction of individual neurons. Elife 5, e10566. doi: 10.7554/eLife.10566.

Elowitz, M.B., Levine, A.J., Siggia, E.D., and Swain, P.S. (2002). Stochastic gene expression in a single cell. Science 297(5584), 1183–1186. doi: 10.1126/science.1070919.

Friedmann, D., Pun, A., Adams, E.L., Lui, J.H., Kebschull, J.M., Grutzner, S.M., et al. (2020). Mapping mesoscale axonal projections in the mouse brain using a 3D convolutional network. Proc Natl Acad Sci U S A 117(20), 11068–11075. doi: 10.1073/pnas.1918465117.

Gagnon, D., and Parent, M. (2014). Distribution of VGLUT3 in highly collateralized axons from the rat dorsal raphe nucleus as revealed by single-neuron reconstructions. PLoS One 9(2), e87709. doi: 10.1371/journal.pone.0087709.

Gatto, R. (2013). The von Mises-Fisher distribution of the first exit point from the hypersphere of the drifted Brownian motion and the density of the first exit time. Stat Prob Lett 83, 1669–1676.

Gershon, M.D. (2009). Enteric serotonergic neurones … finally! J Physiol 587(3), 507. doi: 10.1113/jphysiol.2008.167676.

Gershon, M.D., and Tack, J. (2007). The serotonin signaling system: from basic understanding to drug development for functional GI disorders. Gastroenterology 132(1), 397–414. doi: 10.1053/j.gastro.2006.11.002.

Gu, C. (2021). Rapid and reversible development of axonal varicosities: A new form of neural plasticity. Front Mol Neurosci 14, 610857. doi: 10.3389/fnmol.2021.610857.

Hatada, Y., Wu, F., Silverman, R., Schacher, S., and Goldberg, D.J. (1999). *En passant* synaptic varicosities form directly from growth cones by transient cessation of growth cone advance but not of actin-based motility. J Neurobiol 41(2), 242–251.

Hawthorne, A.L., Wylie, C.J., Landmesser, L.T., Deneris, E.S., and Silver, J. (2010). Serotonergic neurons migrate radially through the neuroepithelium by dynamin-mediated somal translocation. J Neurosci 30(2), 420–430. doi: 10.1523/jneurosci.2333-09.2010.

Hellwig, B., Schüz, A., and Aertsen, A. (1994). Synapses on axon collaterals of pyramidal cells are spaced at random intervals: A Golgi study in the mouse cerebral cortex. Biol Cybern 71(1), 1–12. doi: 10.1007/bf00198906.

Hendricks, T., Francis, N., Fyodorov, D., and Deneris, E.S. (1999). The ETS domain factor *Pet-1* is an early and precise marker of central serotonin neurons and interacts with a conserved element in serotonergic genes. J Neurosci 19(23), 10348–10356.

Hingorani, M., Viviani, A.M.L., Sanfilippo, J.E., and Janušonis, S. (2022). High-resolution spatiotemporal analysis of single serotonergic axons in an *in vitro* system. Front Neurosci 16, 994735. doi: 10.3389/fnins.2022.994735.

Hoff, P.D. (2009). Simulation of the matrix Bingham-von Mises-Fisher distribution, with applications to multivariate and relational data. J. Comput. Graph. Stat. 18(2), 438–456.

Hornung, J.P. (2003). The human raphe nuclei and the serotonergic system. J Chem Neuroanat 26(4), 331–343. doi: 10.1016/j.jchemneu.2003.10.002.

Huang, C.X., Zhao, Y., Mao, J., Wang, Z., Xu, L., Cheng, J., et al. (2021). An injury-induced serotonergic neuron subpopulation contributes to axon regrowth and function restoration after spinal cord injury in zebrafish. Nat Commun 12(1), 7093. doi: 10.1038/s41467-021-27419-w.

Jacobs, B.L., and Azmitia, E.C. (1992). Structure and function of the brain serotonin system. Physiol Rev 72(1), 165–229. doi: 10.1152/physrev.1992.72.1.165.

Janušonis, S. (2018). Some galeomorph sharks express a mammalian microglia-specific protein in radial ependymoglia of the telencephalon. Brain Behav Evol 91(1), 17–30. doi: 10.1159/000484196.

Janušonis, S., and Detering, N. (2019). A stochastic approach to serotonergic fibers in mental disorders. Biochimie 161, 15–22. doi: 10.1016/j.biochi.2018.07.014.

Janušonis, S., Detering, N., Metzler, R., and Vojta, T. (2020). Serotonergic axons as fractional Brownian motion paths: Insights into the self-organization of regional densities. Front Comput Neurosci 14, 56. doi: 10.3389/fncom.2020.00056.

Janušonis, S., Haiman, J.H., Metzler, R., and Vojta, T. (2023). Predicting the distribution of serotonergic axons: A supercomputing simulation of reflected fractional Brownian motion in a 3D-mouse brain model. Front Comput Neurosci 17, 1189853. doi: 10.3389/fncom.2023.1189853.

Janušonis, S., Mays, K.C., and Hingorani, M.T. (2019). Serotonergic axons as 3D-walks. ACS Chem Neurosci 10(7), 3064–3067. doi: 10.1021/acschemneuro.8b00667.

Jin, Y., Dougherty, S.E., Wood, K., Sun, L., Cudmore, R.H., Abdalla, A., et al. (2016). Regrowth of serotonin axons in the adult mouse brain following injury. Neuron 91(4), 748–762. doi: 10.1016/j.neuron.2016.07.024.

Kajstura, T.J., Dougherty, S.E., and Linden, D.J. (2018). Serotonin axons in the neocortex of the adult female mouse regrow after traumatic brain injury. J Neurosci Res 96(4), 512–526. doi: 10.1002/jnr.24059.

Kitt, M.M., Tabuchi, N., Spencer, W.C., Robinson, H.L., Zhang, X.L., Eastman, B.A., et al. (2022). An adult-stage transcriptional program for survival of serotonergic connectivity. Cell Rep 39(3), 110711. doi: 10.1016/j.celrep.2022.110711.

Lee, C., Zhang, Z., and Janušonis, S. (2022). Brain serotonergic fibers suggest anomalous diffusion-based dropout in artificial neural networks. Front Neurosci 16, 949934. doi: 10.3389/fnins.2022.949934.

Lesch, K.P., and Waider, J. (2012). Serotonin in the modulation of neural plasticity and networks: implications for neurodevelopmental disorders. Neuron 76(1), 175–191. doi: 10.1016/j.neuron.2012.09.013.

Lidov, H.G., and Molliver, M.E. (1982). Immunohistochemical study of the development of serotonergic neurons in the rat CNS. Brain Res Bull 9(1-6), 559–604. doi: 10.1016/0361-9230(82)90164-2.

Lillesaar, C. (2011). The serotonergic system in fish. J Chem Neuroanat 41(4), 294–308. doi: 10.1016/j.jchemneu.2011.05.009.

López, J.M., and González, A. (2014). Organization of the serotonergic system in the central nervous system of two basal actinopterygian fishes: the Cladistians *Polypterus senegalus* and *Erpetoichthys calabaricus*. Brain Behav Evol 83(1), 54–76. doi: 10.1159/000358266.

Luchetti, A., Bota, A., Weitemier, A., Mizuta, K., Sato, M., Islam, T., et al. (2020). Two functionally distinct serotonergic projections into hippocampus. J Neurosci. 40(25), 4936–4944. doi: 10.1523/jneurosci.2724-19.2020.

Ma, D., Deng, B., Sun, C., McComb, D.W., and Gu, C. (2022). The mechanical microenvironment regulates axon diameters visualized by cryo-electron tomography. Cells 11(16), 2533. doi: 10.3390/cells11162533.

Maddaloni, G., Bertero, A., Pratelli, M., Barsotti, N., Boonstra, A., Giorgi, A., et al. (2017). Development of serotonergic fibers in the post-natal mouse brain. Front Cell Neurosci 11, 202. doi: 10.3389/fncel.2017.00202.

Maia, G.H., Soares, J.I., Almeida, S.G., Leite, J.M., Baptista, H.X., Lukoyanova, A.N., et al. (2019). Altered serotonin innervation in the rat epileptic brain. Brain Res Bull 152, 95–106. doi: 10.1016/j.brainresbull.2019.07.009.

Mardia, K.V., and Jupp, P.E. (2000). Directional Statistics. New York: John Wiley & Sons.

Morgan, A.A., Alves, N.D., Stevens, G.S., Yeasmin, T.T., Mackay, A., Power, S., et al. (2023). Medial prefrontal cortex serotonin input regulates cognitive flexibility in mice. bioRxiv. doi: 10.1101/2023.03.30.534775.

Muzumdar, M.D., Tasic, B., Miyamichi, K., Li, L., and Luo, L. (2007). A global double-fluorescent Cre reporter mouse. Genesis 45(9), 593–605. doi: 10.1002/dvg.20335.

Navabpour, S., Kwapis, J.L., and Jarome, T.J. (2020). A neuroscientist’s guide to transgenic mice and other genetic tools. Neurosci Biobehav Rev 108, 732–748. doi: 10.1016/j.neubiorev.2019.12.013.

Neal, K.B., Parry, L.J., and Bornstein, J.C. (2009). Strain-specific genetics, anatomy and function of enteric neural serotonergic pathways in inbred mice. J Physiol 587(3), 567–586. doi: 10.1113/jphysiol.2008.160416.

Okaty, B.W., Commons, K.G., and Dymecki, S.M. (2019). Embracing diversity in the 5-HT neuronal system. Nat Rev Neurosci 20(7), 397–424. doi: 10.1038/s41583-019-0151-3.

Okaty, B.W., Sturrock, N., Escobedo Lozoya, Y., Chang, Y., Senft, R.A., Lyon, K.A., et al. (2020). A single-cell transcriptomic and anatomic atlas of mouse dorsal raphe *Pet1* neurons. Elife 9. doi: 10.7554/eLife.55523.

Paul, E.R., Schwieler, L., Erhardt, S., Boda, S., Trepci, A., Kämpe, R., et al. (2022). Peripheral and central kynurenine pathway abnormalities in major depression. Brain Behav Immun 101, 136–145. doi: 10.1016/j.bbi.2022.01.002.

Pewsey, A., and Garcia-Portugués, E. (2021). Recet advances in directional statistics. Test 30(3), 1–58. doi: 10.1007/s11749-021-00759-x.

Pratelli, M., Migliarini, S., Pelosi, B., Napolitano, F., Usiello, A., and Pasqualetti, M. (2017). Perturbation of serotonin homeostasis during adulthood affects serotonergic neuronal circuitry. eNeuro 4(2). doi: 10.1523/eneuro.0376-16.2017.

Raj, A., and van Oudenaarden, A. (2008). Nature, nurture, or chance: stochastic gene expression and its consequences. Cell 135(2), 216–226. doi: 10.1016/j.cell.2008.09.050.

Ren, J., Friedmann, D., Xiong, J., Liu, C.D., Ferguson, B.R., Weerakkody, T., et al. (2018). Anatomically defined and functionally distinct dorsal raphe serotonin sub-systems. Cell 175(2), 472–487.e420. doi: 10.1016/j.cell.2018.07.043.

Ren, J., Isakova, A., Friedmann, D., Zeng, J., Grutzner, S.M., Pun, A., et al. (2019). Single-cell transcriptomes and whole-brain projections of serotonin neurons in the mouse dorsal and median raphe nuclei. Elife 8. doi: 10.7554/eLife.49424.

Romanczuk, P., Bär, M., Ebeling, W., and Lindner, B. (2012). Active Brownian particles. Eur. Phys. J. Special Topics 202, 1–162.

Saito, T. (2023). Langevin analogy brtween particle trajectories and polymer configurations. Phys Rev E 107, 034502.

Sako, H., Kojima, T., and Okado, N. (1986). Immunohistochemical study on the development of serotoninergic neurons in the chick: I. Distribution of cell bodies and fibers in the brain. J Comp Neurol 253(1), 61–78. doi: 10.1002/cne.902530106.

Shaw, G., and Bray, D. (1977). Movement and extension of isolated growth cones. Exp Cell Res 104(1), 55–62. doi: 10.1016/0014-4827(77)90068-4.

Shepherd, G.M., Raastad, M., and Andersen, P. (2002). General and variable features of varicosity spacing along unmyelinated axons in the hippocampus and cerebellum. Proc Natl Acad Sci U S A 99(9), 6340–6345. doi: 10.1073/pnas.052151299.

Stuesse, S.L., Stuesse, D.C., and Cruce, W.L. (1995). Raphe nuclei in three cartilaginous fishes, *Hydrolagus colliei*, *Heterodontus francisci*, and *Squalus acanthias*. J Comp Neurol 358(3), 414–427. doi: 10.1002/cne.903580308.

Tanabe, A., Fukumizu, K., Oba, S., T., T., and S., I. (2007). Parameter estimation for von Mises-Fisher distributions. Comput. Stat. 22, 145–157. doi: 10.1007/s00180-007-0030-7.

Teissier, A., Soiza-Reilly, M., and Gaspar, P. (2017). Refining the role of 5-HT in postnatal development of brain circuits. Front Cell Neurosci 11, 139. doi: 10.3389/fncel.2017.00139.

Tsagris, M., Athineou, G., C., A., A., S., E., A., and M.J., W. (2023). Package ‘Directional’: A collection of functions for directional data analysis. CRAN cran.r-project.org/web/packages/Directional/.

Ueda, S., Nojyo, Y., and Sano, Y. (1984). Immunohistochemical demonstration of the serotonin neuron system in the central nervous system of the bullfrog, Rana catesbeiana. Anat Embryol (Berl) 169(3), 219–229. doi: 10.1007/bf00315627.

Vargas, M.V., Dunlap, L.E., Dong, C., Carter, S.J., Tombari, R.J., Jami, S.A., et al. (2023). Psychedelics promote neuroplasticity through the activation of intracellular 5-HT_2A_ receptors. Science 379(6633), 700–706. doi: 10.1126/science.adf0435.

Vojta, T., Halladay, S., Skinner, S., Janušonis, S., Guggenberger, T., and Metzler, R. (2020). Reflected fractional Brownian motion in one and higher dimensions. Phys Rev E 102(3-1), 032108. doi: 10.1103/PhysRevE.102.032108.

Wang, W., Balcerek, M., Burnecki, K., Chechkin, A.V., Janušonis, S., Slezak, J., et al. (2023). Memory-multi-fractional Brownian motion with continuous correlations. *arXiv* 2303.01551.

Weber, T., Böhm, G., Hermann, E., Schütz, G., Schönig, K., and Bartsch, D. (2009). Inducible gene manipulations in serotonergic neurons. Front Mol Neurosci 2, 24. doi: 10.3389/neuro.02.024.2009.

Wilson, M.A., and Molliver, M.E. (1991). The organization of serotonergic projections to cerebral cortex in primates: Regional distribution of axon terminals. Neuroscience 44(3), 537–553. doi: 10.1016/0306-4522(91)90076-z.

Wood, A.T.A. (1994). Simulation of the von Mises-Fisher distribution. Commun. Stat. – Simula 23(1), 157–164.

Yabut, J.M., Crane, J.D., Green, A.E., Keating, D.J., Khan, W.I., and Steinberg, G.R. (2019). Emerging roles for serotonin in regulating metabolism: New implications for an ancient molecule. Endocr Rev 40(4), 1092–1107. doi: 10.1210/er.2018-00283.

Yoshinobu, K., Araki, M., Morita, A., Araki, M., Kokuba, S., Nakagata, N., et al. (2021). Tamoxifen feeding method is suitable for efficient conditional knockout. Exp Anim 70(1), 91–100. doi: 10.1538/expanim.19-0138.

Zhang, H., Fujitani, Y., Wright, C.V., and Gannon, M. (2005). Efficient recombination in pancreatic islets by a tamoxifen-inducible Cre-recombinase. Genesis 42(3), 210–217. doi: 10.1002/gene.20137.

